# Evolutionary success of a parasitic B chromosome rests on gene content

**DOI:** 10.1101/683417

**Authors:** Francisco J. Ruiz-Ruano, Beatriz Navarro-Domínguez, María Dolores López-León, Josefa Cabrero, Juan Pedro M. Camacho

## Abstract

Supernumerary (B) chromosomes are dispensable genomic elements found in most kinds of eukaryotic genomes. Many show drive mechanisms that give them an advantage in transmission, but how they achieve it remains a mystery. The recent finding of protein-coding genes in B chromosomes has opened the possibility that their evolutionary success is based on their genetic content. Using a protocol based on mapping genomic DNA Illumina reads from B-carrying and B-lacking individuals on the coding sequences of *de novo* transcriptomes from the same individuals, we identified 25 protein-coding genes in the B chromosome of the migratory locust, 15 of which showed a full coding region. Remarkably, one of these genes (*apc1*) codes for the large subunit of the Anaphase Promoting Complex or Cyclosome (APC/C), an E3 ubiquitin ligase involved in the metaphase-anaphase transition. Sequence comparison of A and B chromosome gene paralogs showed that the latter show B-specific nucleotide changes, neither of which putatively impairs protein function. These nucleotide signatures allowed identifying B-derived transcripts in B-carrying transcriptomes, and demonstrated that they show about similar frequency as A-derived ones. Since B-carrying individuals show higher amounts of *apc1* transcripts than B-lacking ones, the putatively higher amount of APC1 protein might induce a faster metaphase-anaphase transition in spite of orientation of the two B chromosome chromatids towards the same pole during metaphase, thus facilitating B chromosome non-disjunction. Therefore, *apc1* is the first protein-coding gene uncovered in a B chromosome that might be responsible for B chromosome drive.

**Significance Statement:** The genome of the migratory locust harbors a parasitic chromosome that arose about 2 million years ago. It is widespread in natural populations from Asia, Africa, Australia and Europe, i.e. all continents where this species lives. The secret for such an extraordinary evolutionary success is unveiled in this report, as B chromosomes in this species contain active protein-coding genes whose transcripts might interfere with gene expression in the host genome (the A chromosomes), thus facilitating B chromosome mitotic and meiotic drive to provide the transmission advantage which grants its success. One of the B-chromosomal genes (*apc1*) codes for the large subunit of the Anaphase Promoting Complex or Cyclosome (APC/C) whose expression might provide a mechanistic explanation for B chromosome drive.

## Introduction

After 112 years since they were uncovered (1), B chromosomes continue being an enigmatic part of eukaryote genomes. They were first considered as merely genetically inert passengers of eukaryote genomes (2), a view supported by others (3) but criticized by those who argued that B chromosomes are beneficial (4) or parasitic (5) elements. In fact, as in most cases of selfish genetic elements, phenotypic effects of B chromosomes are usually modest (6), but they frequently decrease fertility (7, 8). An extreme case was found in the wasp *Nasonia vitripennis*, where a B chromosome decreases the fitness of the host genome to zero by the elimination of all paternal chromosomes accompanying it in the spermatozoon (9).

Thirteen years ago, two findings suggested that B chromosomes are not inert elements, namely the first molecular evidence of gene activity on B chromosomes (10) and the finding of protein-coding genes in them (11). It was later found evidence for transcription of ribosomal DNA in B chromosomes (12) and that rRNA transcripts from a B chromosome are functional (13), The first evidence for transcription of a protein-coding gene on B chromosomes was provided by Tryfonov et al. (14).

The arrival of next generation sequencing (NGS) has drastically accelerated B chromosome research during last years. The pioneering work by Martis et al. (15) revealed that B chromosomes in rye are rich in gene fragments, some of which are pseudogenical and transcribed (16). Likewise, Valente et al. (17) found that a B chromosome in fish contained thousands of gene-like sequences, most of them being fragmented but a few remaining largely intact, with at least three of them being transcriptionally active. Likewise, Navarro-Domínguez et al. (18) found evidence for the presence of ten protein-coding genes in the B chromosome of the grasshopper *Eyprepocnemis plorans*, five of which were active in B-carrying individuals, including three which were apparently pseudogenic. Taken together, these results indicate that B chromosomes are not as inert as previously thought, since they contain protein-coding genes which are actively transcribed even in the case of being pseudogenic.

However, the extent to which B chromosomes express their genetic content is still unknown. Lin et al. (19) characterized B-chromosome-related transcripts in maize and concluded that the maize B chromosome harbors few transcriptionally active sequences and might influence transcription in A chromosomes. Recently, Huang et al. (20) have found that the expression of maize A chromosome genes is influenced by the presence of B chromosomes, and that four up-regulated genes are actually present in the B chromosomes. Notably, it has been recently shown that rye B chromosomes carry active Argonaute-like genes and that B-encoded AGO4B protein variants show in vitro RNA slicer activity (21), thus going a step further in demonstrating the functionality of B chromosome gene paralogs. Recently, it has been shown that B chromosome presence in *E. plorans* triggers transcriptional changes being consistent with some of the effects previously reported in this species (22).

The genome of the migratory locust (*Locusta migratoria*) harbors a highly successful B chromosome found in most natural populations from Asia (23–26), Africa (27), Australia (28) and Europe (29). The wide geographical range of this B chromosome is based on mitotic and meiotic drive mechanisms in males and females, respectively (30), along with their modest harmful effects (31). Here we use NGS approaches to search for protein-coding genes in B chromosomes with the potential to manipulate cell division, by means of genomic and transcriptomic analyses. We found that the B chromosome of the migratory locust contains at least 25 protein-coding genes. One of them codes for the large subunit of the APC/C protein complex, an E3 ubiquitin ligase involved in the control of metaphase-anaphase progression thus giving a reasonable explanation to the non-disjuntion events characterizing B chromosome drive mechanisms in this species.

## Results

### Searching for protein-coding genes in the B chromosome

*De novo* transcriptome assembly of the 12 RNA libraries from Cádiz (testis and hind leg from six males) yielded 523,445 contigs (N50= 792 nt), 108,517 of which included CDSs higher than 300 nt (N50= 891 nt). After clustering (to reduce redundancy) we got a final reference assembly including 49,476 CDSs (N50= 975 nt). In total, gDNA mappings yielded counts for 47,763 CDS contigs. To discard contigs that might include repetitive DNA (i.e. those with more than 4 copies per genome), or those showing putative assembling problems (i.e. those with less than 0.5 copies), we limited our analysis to the 28,667 CDS contigs showing 0.5-4 copies in the B-lacking (0B) genomes (Fig. *1A*, Dataset S1).

**Figure 1.**
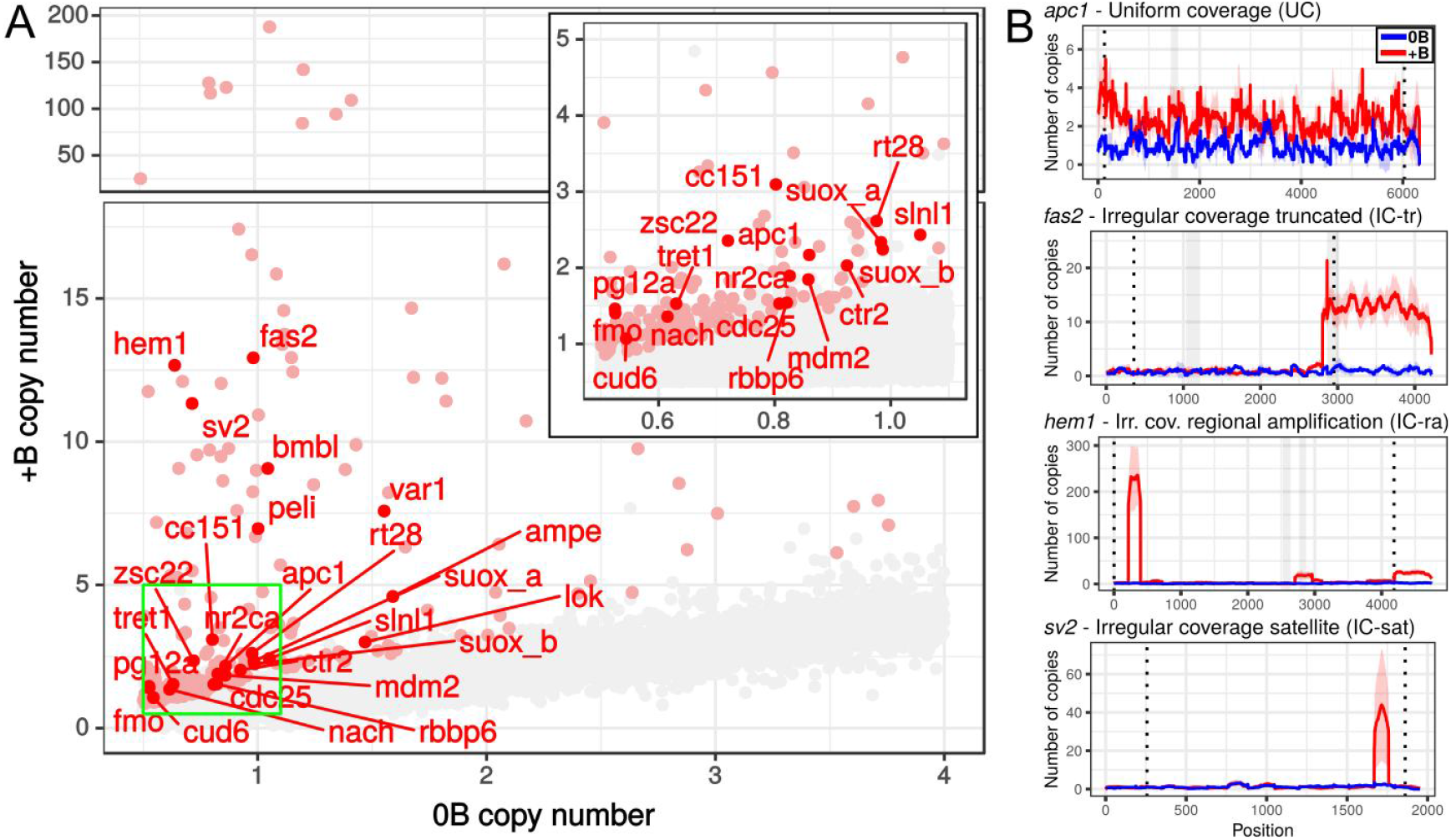
Detection of protein-coding genes residing in the B chromosome of *Locusta migratoria* by mapping gDNA Illumina reads obtained from four B-carrying (+B) and two B-lacking (0B) males on a *de novo* transcriptome built with Illumina reads coming from testis and hind leg RNA libraries from the same individuals (Cádiz population). (A) Scatterplot showing the mean copy number for 27,313 CDSs in 0B and +B males, with enlargement of the region included in the green box (inset). The CDSs that in presence of B chromosomes showed a genomic fold change [gFC= log2(+B/0B)] higher than 0.585 in the four B-carrying individuals (72 contigs in total, see Dataset S1) are shown in light red color, and those failing to reach this threshold are noted in grey color. In addition, the 25 protein-coding genes which were validated by qPCR are noted in dark red color for the contig showing the highest gFC value. (B) Examples of the four coverage patterns found: UC= uniform coverage, IC-tr= Irregular coverage by truncation, IC-ra= Irregular coverage by regional amplification, and IC-sat= Irregular coverage by satellite DNA formation.

After gDNA SSAHA2 mappings and genomic coverage fold change [gFC= log2(+B/0B)] calculations, we selected 271 contigs showing gFC>0.585 in all four B-carrying individuals analysed (Dataset S1). This threshold implies assuming the existence of one gene copy per A or B chromosome. Since B-carrying individuals came from the same population (Cádiz), we also assumed that B chromosomes in different individuals show similar gene content. Finally, 72 of these contigs were annotated as protein-coding genes (Table S1). As several contigs were annotated to the same gene, the final list finally included 61 different genes (Table S2). We thoroughly analyzed the sequence of these 61 genes trying to infer as much of their length as possible, thus including some UTR regions, and manually assembling the sequence of contigs belonging to the same gene. Unfortunately, we could not use the published genome of *L. migratoria* (32) for this purpose because, in this assembly, many genes are found fragmented in several scaffolds and, frequently, the predicted CDS is incomplete. Alternatively, our *de novo* 0B transcriptome provided information to extend the known sequence of the 61 genes. As these sequences were obtained using exclusively B-lacking individuals, we can conclude that they correspond to sequences in the A chromosomes.

The sequences of these 61 genes were used as reference to separately map the reads from each of all B-carrying and B-lacking gDNA libraries from Cádiz, to examine coverage variation per nucleotide along CDS+UTRs length (Table S2). Log2 of the quotient between the coefficients of variation of +B and 0B individuals indicated a fold change in coverage variation along gene sequence (cvFC). We interpreted that negative values of this parameter indicated uniform coverage (UC) in the B-chromosome paralog whereas positive values indicated irregular coverage (IC) (40 and 21 genes, respectively) (Table S2). The latter type can be subdivided into three subtypes: those being truncated (IC-tr genes), those showing regional amplification (IC-ra) and those showing high coverage only for a short region being tandemly repeated in the B-carrying gDNA, thus yielding a satellite DNA (IC-sat) (see examples in Fig. 1*B*).

We arranged the 61 genes in order of decreasing gFC values (in the high coverage region, in case of IC genes), 30 of which showed gFC>0.585 in all four B-carrying individuals analyzed. We then performed qPCR validation for 15 out of these 30 genes, preferentially choosing those with UC pattern and functions related with cell cycle. Remarkably, qPCR showed that 14 of these 15 genes showed overabundance in the B-carrying gDNAs, and 12 of them showed gFC>1 suggesting that all 22 genes surpassing this gFC value are present in the B chromosome (Table S2). The exception was the *skp2* gene, with gFC= 0.929, for which reason we added other genes with gFC values lower than this, to the B-genes list, only if they had been qPCR validated (i.e. *cdc25* and *cud6*, see Table S2). We also analyzed, by qPCR, five genes passing the gFC>0.585 threshold in only three individuals, and five genes showing gFC<0.585 as negative control (Table S2, Fig. S1). Whereas all latter genes failed to be validated, one out of the five former genes (*mdm2*) was validated by qPCR, and it was added to the list of B-genes thus summing up 25 B-genes (Tables 1 and S2). In total, we found 15 genes showing the UC coverage pattern and 10 showing IC patterns (5 IC-tr, 4 IC-ra and 1 IC-sat). Obviously, total coverage and methodological limitations of our experiments indicate that the list of B-genes is not yet complete.

**Table 1.**
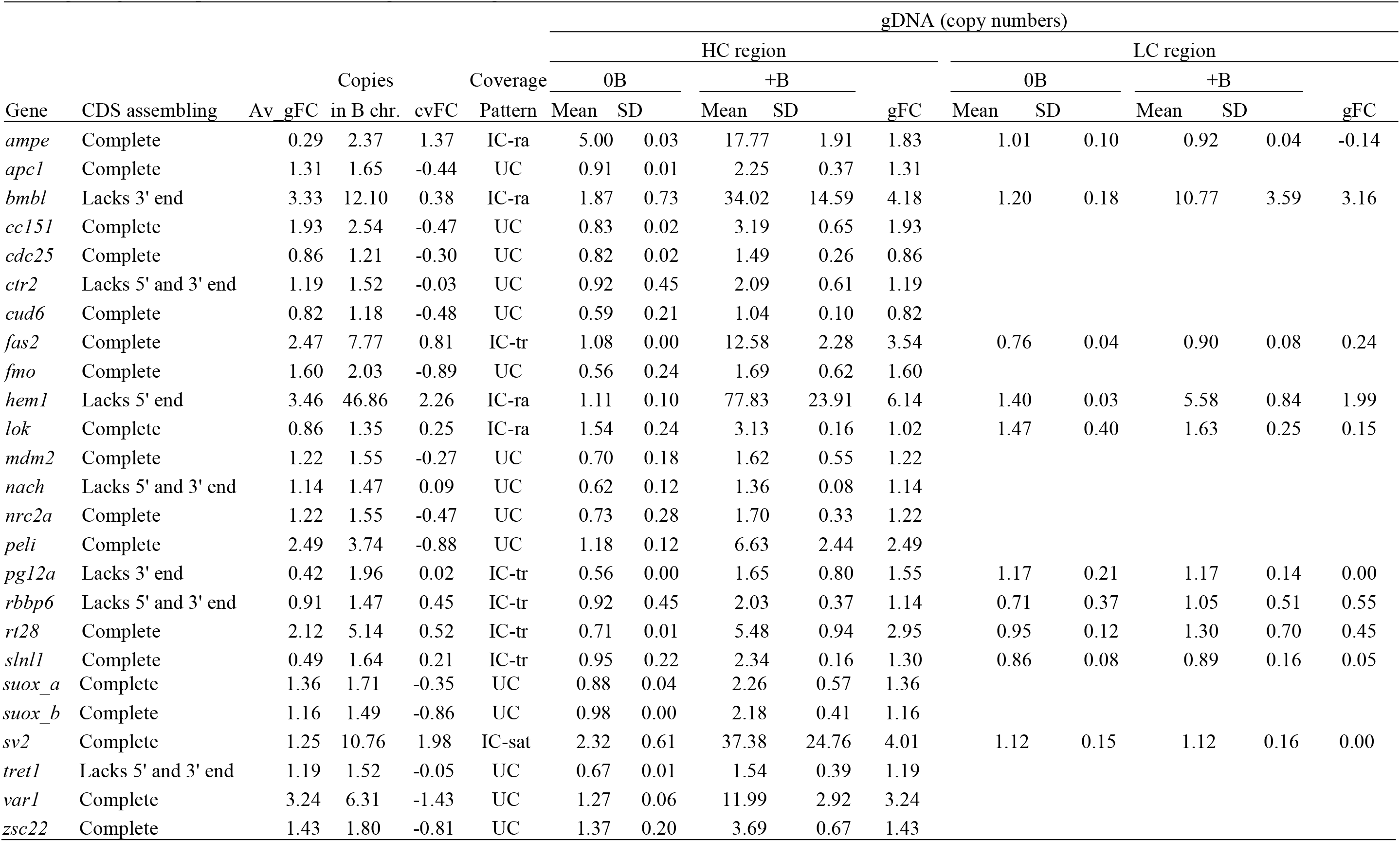
Basical structural features of the 25 B-chromosome genes. Average genomic fold change (Av_gFC) vas calculated as log2 the +B/0B quotient between the mean nucleotidic coverages along CDS length. The distribution pattern of coverage along CDS length was measured by log2 the +B/0B quotient of the coefficients of variation along CDS length (cvFC). Negative values of cvFC indicated uniform coverage suggesting that the CDS is complete in the B chromosome (UC pattern). Positive values indicated irregular patterns of coverage (IC). HC= High coverage region. LC= Low coverage region. IC-tr= Irregular Coverage - truncated. IC-ra= Irregular Coverage - regional amplification. IC-sat= Irregular Coverage - satellite.

As an additional test of the reliability of this result, we performed gFC analysis on two Illumina DNA libraries obtained from 0B and +B individuals collected at Padul. Although these samples did not allow applying the same criteria employed above, due to the absence of replicates, the gFC values calculated from them showed very high positive correlation with the average gFC values for the four B-carrying males from Cádiz (Fig. 2*A* and Table S3). In fact, 19 genes in the Padul libraries showed gFC≥1.45 and only two showed it lower than 0.585. These 25 protein-coding genes showed a wide variety of KOG functions (Table 2) presumably reflecting the A chromosome region(s) which gave rise to the B chromosome and probably saying little, as a whole, about B chromosome functions. The whole pathway for gene search is summarized in Fig. S2.

**Table 2.** KOG annotation for the genes found in the B chromosome. The *cc151, ctr2, cud6, nrc2a, pg12a, rbbp6, slnl1* and *tret1* genes were not annotated.

| Gene | Hit | Class | Class description |
| --- | --- | --- | --- |
| *ampe* | KOG1046 | E | Amino acid transport and metabolism |
|  |  | O | Posttranslational modification, protein turnover, chaperones |
| *apc1* | KOG1858 | D | Cell cycle control, cell division, chromosome partitioning |
|  |  | O | Posttranslational modification, protein turnover, chaperones |
| *bmb1* | KOG4673 | K | Transcription |
| *cdc25* | KOG3772 | D | Cell cycle control, cell division, chromosome partitioning |
| *fas2* | KOG3513 | T | Signal transduction mechanisms |
| *fmo* | KOG1399 | Q | Secondary metabolites biosynthesis, transport and catabolism |
| *hem1* | KOG3513 | T | Signal transduction mechanisms |
| *lok* | KOG0615 | D | Cell cycle control, cell division, chromosome partitioning |
| *mdm2* | KOG4172 | O | Posttranslational modification, protein turnover, chaperones |
| *nach* | KOG4294 | P | Inorganic ion transport and metabolism |
|  |  | T | Signal transduction mechanisms |
| *pel1* | KOG3842 | T | Signal transduction mechanisms |
| *rt28* | KOG4078 | J | Translation, ribosomal structure and biogenesis |
| *suox_a* | KOG0535 | C | Energy production and conversion |
| *suox_b* | KOG0535 | C | Energy production and conversion |
|  | KOG4576 | C | Energy production and conversion |
| *sv2* | KOG0253 | R | General function prediction only |
|  | KOG0255 | R | General function prediction only |
| *var1* | KOG1445 | Z | Cytoskeleton |
| *zsc22* | KOG2462 | K | Transcription |

**Figure 2.**
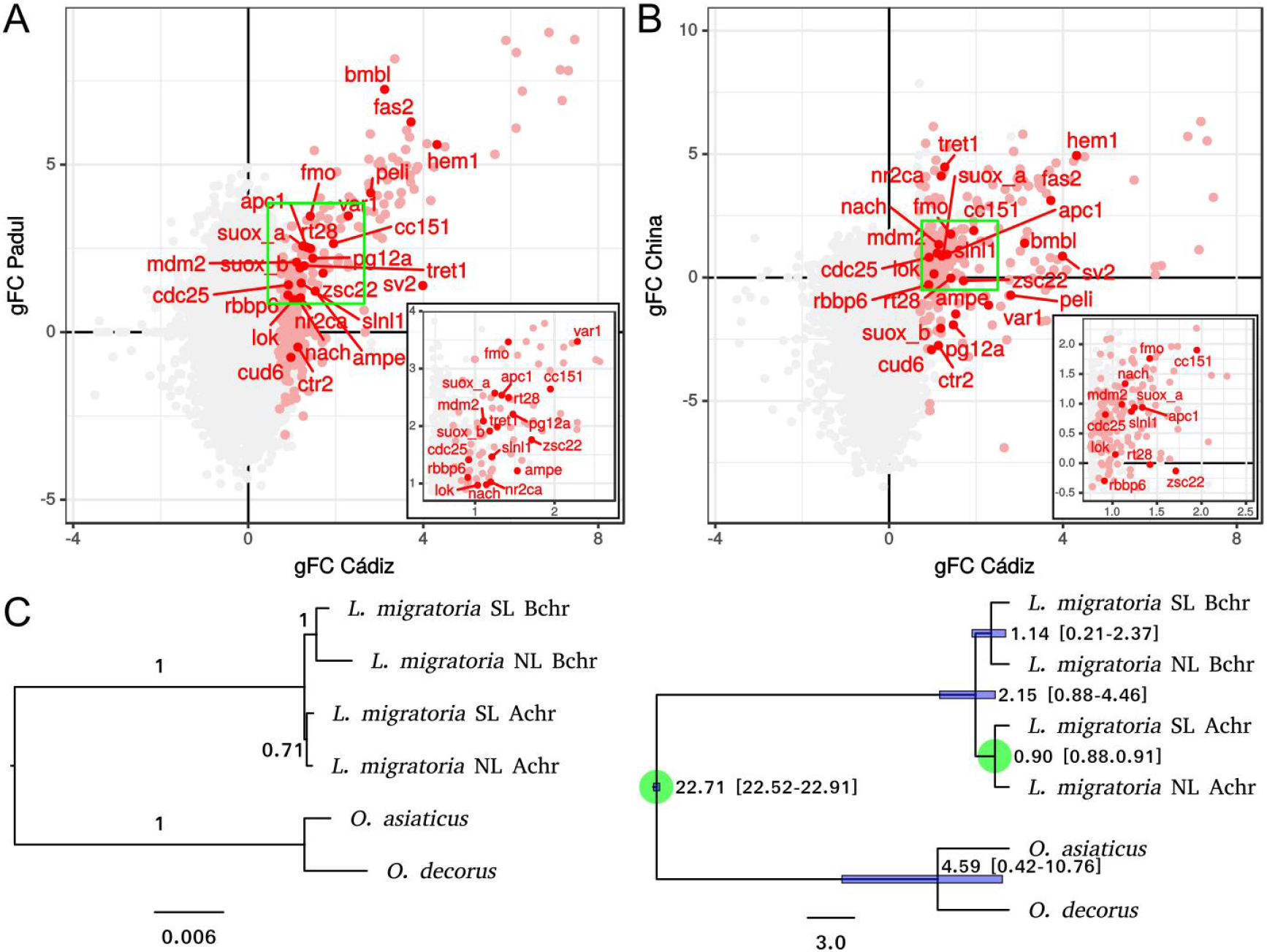
Protein-coding gene content of the B chromosome is highly similar among populations. Color codes in *A* and *B* are as in Figure 1. (*A*) Comparison of gFC values between two Spanish populations at Cádiz and Padul (Granada). The gFC values found for the 23 B chromosome genes found in both populations were positively correlated (rs= 0.88, P<0.000001). (*B*) Comparison of gDNA FC between Cádiz and China libraries. Again, gFC values for the 25 B chromosome genes found in both populations were positively correlated (rs= 0.65, P= 0.00043). (*C*) Phylogram and chronogram built with a 17,411 nt sequence obtained by concatenating ten B-chromosome genes showing B-specific variation. Note that the B chromosome sequences (Bchr) cluster together and are clearly separated from A chromosome ones (Achr). The phylogram on the left shows longer branches for paralogs in B chromosome genes, suggesting accumulation of more changes in B than A gene paralogs. The chronogram on the right suggests that the B chromosome arose in *L. migratoria* about 2 mya, prior to the separation of NL and SL lineages.

### B-specific sequence variation

To ascertain whether B-chromosome genes carry sequence signatures, we searched for DNA sequence variation being specific to B-carrying individuals and thus to the B chromosome gene paralogs. Specifically, we searched for nucleotide variations being present in all four B-carrying gDNA libraries from Cádiz but absent in the gDNA and RNA libraries from the two 0B males analyzed from this same population. This yielded 141 single nucleotide polymorphisms (SNPs) representing B-specific variation (Alt variant) in 19 B-genes (Tables 3 and S4), 79 of which were found in CDS regions whereas the remaining 62 were in UTR regions. This high number of SNPs suggests that B chromosomes in *L. migratoria* are quite old and their DNA sequences have been independently evolving for long from those in the A chromosomes. Remarkably, the genes showing IC pattern, as a whole, did not differ from those showing UC pattern for the number of synonymous (dS) or non-synonymous (dN) substitutions in respect to the A chromosome genes or divergence per nucleotide site (ps) in the CDS (Mann-Whitney test: P>0.05 in all cases). This is probably due to the high variation between genes for these four parameters, as there exists two types of genes in the B chromosome, 13 showing 1-8 synonymous changes, in respect to the A-chromosome paralogs, and 9 showing no change at all (Table 3).

**Table 3.** Nucleotidic variation found in the B chromosome gene paralogs. To score the number of variants, we distinguished between substitutions (s), deletions (d) or insertions (i) in respect to the A chromosome paralogs. We considered deletions and insertions as a single mutation event irrespectively whether they involved 1-14 nt. dS= number of synonymous substitutions; dN= number of non-synonymous substitutions; ps= proportion of variable sites.

| Gene | CDS | Number of variants |  |  |  | dS | dN | dN/dS | ps |
| --- | --- | --- | --- | --- | --- | --- | --- | --- | --- |
|  | length | Total | In 5'-UTR | In 3'-UTR | In CDS |  |  |  |  |
| *ampe* | 2577 | 3s | 1s | 0 | 2s | 0 | 2 |  | 0.0008 |
| *apc1* | 5907 | 12s | 1s | 2s | 9s | 8 | 1 | 0.13 | 0.0015 |
| *bmb1* | 1341 | 13s | 0 | 0 | 13s | 5 | 8 | 1.60 | 0.0097 |
| *cc151* | 1650 | 0 | 0 | 0 | 0 | 0 | 0 |  | 0.0000 |
| *cdc25* | 1950 | 1s | 0 | 1s | 0 | 0 | 0 |  | 0.0000 |
| *ctr2* | 332 | 0 | 0 | 0 | 0 | 0 | 0 |  | 0.0000 |
| *cud6* | 303 | 0 | 0 | 0 | 0 | 0 | 0 |  | 0.0000 |
| *fas2* | 2595 | 10s+2d | 0 | 6s+1d | 4s+1d | 3 | 2 | 0.67 | 0.0019 |
| *fmo* | 1350 | 1s | 0 | 0 | 1s | 0 | 1 |  | 0.0007 |
| *hem1* | 4185 | 24s+1d | 0 | 1s | 23s+1d | 6 | 18 | 3.00 | 0.0057 |
| *lok* | 1425 | 2s | 0 | 1s | 1s | 1 | 0 |  | 0.0007 |
| *mdm2* | 1656 | 0 | 0 | 0 | 0 | 0 | 0 |  | 0.0000 |
| *nach* | 306 | 0 | 0 | 0 | 0 | 0 | 0 |  | 0.0000 |
| *nrc2a* | 420 | 1s | 0 | 0 | 1s | 1 | 0 |  | 0.0024 |
| *pel1* | 1206 | 3s+1d | 0 | 3s+1d | 0 | 0 | 0 |  | 0.0000 |
| *pg12a* | 461 | 0 | 0 | 0 | 0 | 0 | 0 |  | 0.0000 |
| *rbbp6* | 337 | 1s | 0 | 0 | 1s | 0 | 1 |  | 0.0030 |
| *rt28* | 543 | 1s | 0 | 1s | 0 | 0 | 0 |  | 0.0000 |
| *sln1* | 1059 | 8s | 0 | 6s | 2s | 1 | 1 | 1.00 | 0.0019 |
| *suox_a* | 777 | 28s+2i+1d | 4s+2i | 16s+1d | 8s | 1 | 7 | 7.00 | 0.0103 |
| *suox_b* | 1689 | 1s | 0 | 0 | 1s | 1 | 0 |  | 0.0006 |
| *sv2* | 1605 | 5s | 3s | 0 | 2s | 1 | 1 | 1.00 | 0.0012 |
| *tret1* | 1121 | 2s | 0 | 0 | 2s | 1 | 1 | 1.00 | 0.0018 |
| *var1* | 1596 | 4s+1d | 1s | 1s+1d | 2s | 2 | 0 |  | 0.0013 |
| *zsc22* | 633 | 10s+3d | 1s+1d | 6s | 3s+2d | 3 | 2 | 0.67 | 0.0079 |
| Total |  | 141 | 14 | 48 | 79 |  |  |  |  |

### The sequenced genome of L. migratoria contains B-specific variation

The presence of a TE chimera restricted to B chromosomes in Spanish populations (33) in the genome assembly published by Wang et al. (32) suggested that the Chinese female used by these latter authors contained B chromosomes. To get additional insights on this possibility, we analyzed coverage for the 25 B-genes in the gDNA of this female and found significant positive correlation between these gFC values and those found in the Cadiz libraries (Fig. 2*B* and Table S3). In fact, 20 out of the 25 genes showed gFC> 0.585 (gFC from 0.92 to 4.11), whereas five genes showed negative gFC values indicating they are not present in the B chromosome in Chinese *L. migratoria.* Remarkably, two of these genes (*ctr2* and *cud6*) showed also negative gFC values in the Padul gDNA libraries, indicating that these genes might represent recent incorporations to the B chromosome in Cádiz. This conclusion is also supported by their lack of synonymous changes in Cádiz libraries.

We searched for possible evidence of linkage between some of the 25 B-genes in the assembled genome using BLAST (34). Due to the draft state of this genome, we only could assess linkage for three of these genes (*slnl1, suox_a* and *suox_b*). The fact that the scaffold no. 43118 ended with the first exon of the *suox_a* gene and scaffold no. 894 began with the second exon of this same gene, allowed us to infer the contiguity of these two scaffolds and the linkage of these three genes (Fig. S3).

The fact that many IC-tr and IC-ra genes showed exactly the same HC and LC regions in the libraries from Spain and China (for instance, see coverage profiles for *fas2* in Dataset S2) could not be explained by chance. Finally, we searched for the presence of the B-specific SNPs in the gDNA library of the Chinese female, and found that 84% of Alt sequence variants (i.e. 119 out of the 141 found in Cádiz libraries) were also present in the Chinese library. Additionally, we searched for B chromosome sequence signatures in 16 transcriptome bioprojects available in SRA, performed on individuals from Asia (Table S5). We performed SNP calling for the 141 SNPs previously found (Table S6) and verified the presence of Alt SNPs in all of them (average= 93 SNPs, minimum= 47, maximum= 125). Taken together, our results strongly suggest that the *L. migratoria* genome published by Wang et al. (32) included B chromosome sequences (Table S4).

### B chromosome age estimation

A concatenated alignment of the coding sequences for 10 B-genes found in A and B chromosomes in Spanish and Chinese *L. migratoria*, and those found in the transcriptomes of *O. decorus* and *O. asiaticus*, showed a total length of 17,411 nt. A Neighbor-Joining phylogram built with this alignment revealed that A and B chromosome sequences were in separate clusters, with B sequences showing longer branches suggesting higher accumulation of nucleotide changes in the B chromosome (Fig. 2*C*). A chronogram built by BEAST placed the origin of B chromosome sequences (and thus B chromosomes) 2.15 Mya, with 95% highest posterior density (HPD) interval of 0.88-4.46 Mya (Fig. 2*C*). Remarkably, B chromosomes in *L. migratoria* arose prior to the separation of the Northern and Southern lineages described in this species by Ma et al. (35), which explains B presence in both lineages.

### Activity of B chromosome gene paralogs

A first indication of gene transcription in the B chromosome was given by the fold change observed in testis and hind leg transcriptomes from Cádiz [tFC= log2(+B/0B) RNA], showing that most contigs being over-represented in the B-carrying gDNA libraries were also over-represented in the transcriptomes of the same individuals (Figs. 3*A* and 3*B*). In fact, 12 out of the 25 B-genes showed tFC>1 in testis or leg transcriptomes (Table S3).

**Figure 3.**
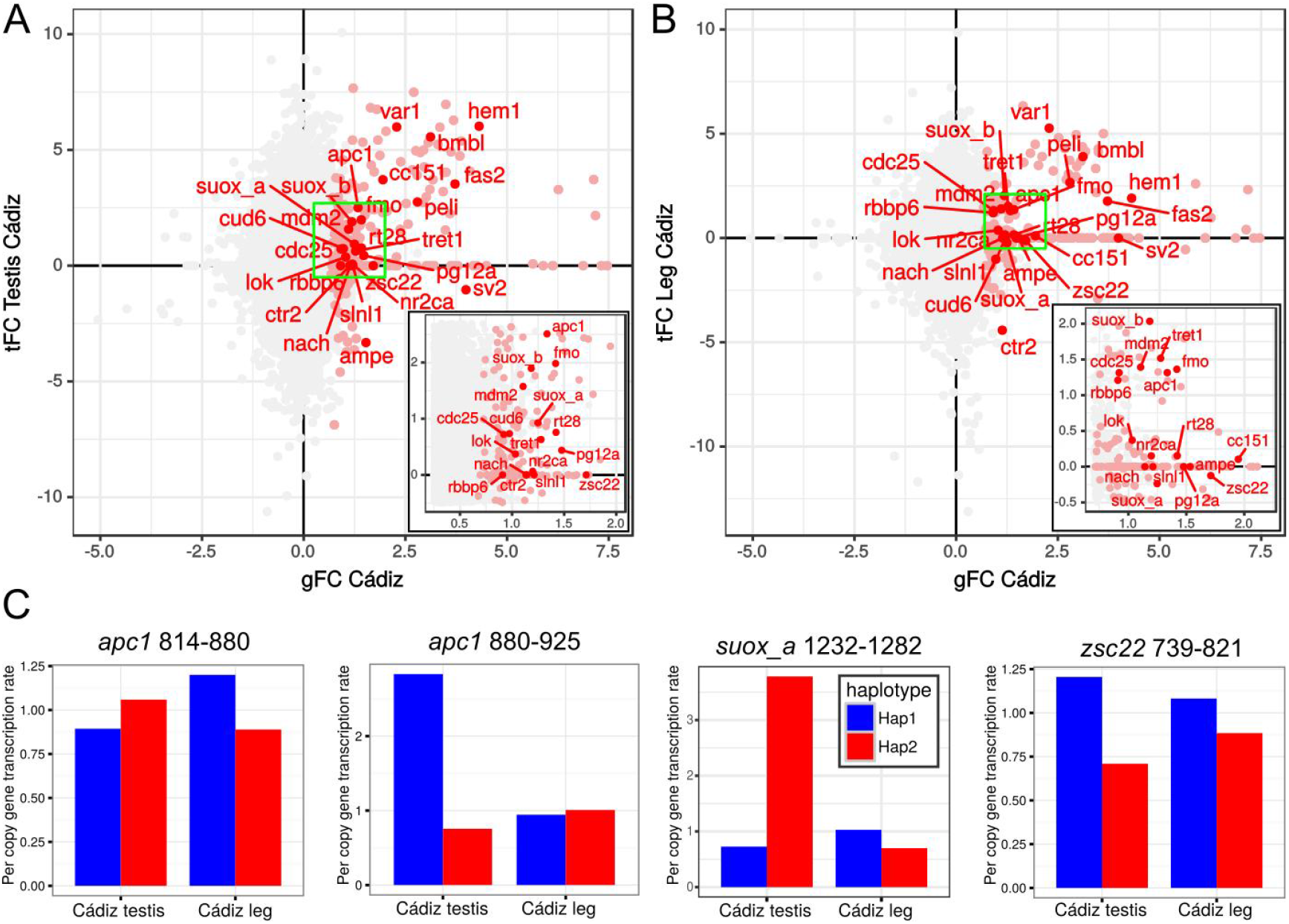
Transcription of B chromosome genes in testis and hind legs of *L. migratoria* males. (*A* and *B*) Transcriptional fold change [(tFC= log2(+B/0B)] observed in B-carrying RNA libraries (Y-axis) in respect to gFC in testis (*A*) and leg (*B*). Color codes in *A* and *B* are as in Figure 1. The green box is enlarged in the inset. Note that many B-genes showed tFC≥1. (*C*) Per copy gen transcription rate (GTR) of B-genes in B-carrying individuals from Cádiz, calculated as the quotient between the numbers of Illumina reads carrying a given haplotype in RNA and gDNA from a same individual. Haplotypes were defined by two (*apc1*), three (*suox_a*) or four (*zsc22*) B-specific SNPs located close enough to be visualized in a same Illumina read. The numbers following gene name indicate the nucleotide positions of the two more distant SNPs defining the haplotype. In all cases, Hap1 (blue bar) was the only haplotype found in 0B individuals and Hap2 (red bar) was exclusive of B-carrying individuals. Note that the A- and B-chromosome haplotypes showed about similar TI.

Estimations of per copy gene transcription rate (GTR) of B-specific gene paralogs, by scoring Alt haplotypes directly in the Illumina reads, revealed that seven out of the ten genes analyzed showed GTR>50% in testis, with only one gene showing no transcription at all and, remarkably, it shows the IC-sat pattern and it is thus likely that the B chromosome carries a satellite arisen from this gene but not the gene itself. Remarkably, in the leg transcriptomes, five genes showed GTR>50% and four genes showed no expression (Tables S7 and S8). Taken together, these results indicate that the B chromosome is not completely silenced but, on the contrary, shows high levels of transcription, especially in testis (Fig. 3*C*).

## Discussion

We uncover here the presence of 25 protein-coding genes in the B chromosome of *L. migratoria*. Of course, this is by no means the complete set of B chromosome genes, and future research with higher coverage will surely uncover some more. As summarized in Table 1, 15 of these genes appear to include a full coding region since they showed uniform coverage (UC pattern). 11 of these genes (*apc1, cc151, cdc25, fmo, mdm2, peli, suox_a, suox_b, tret1, var1* and *zsc22*) showed up-regulation in B-carrying males (tFC≥1 in testis and/or leg transcriptomes) (see Tables 1 and S3). The finding of SNPs being exclusive of B-carrying individuals in five of these UC-genes (*apc1, peli, suox_a, var1* and *zsc22*) allowed defining haplotypes carrying B-specific signatures, and all of them were detected at high proportion in the +B Illumina RNA libraries. We interpret these results as evidence that protein-coding gene paralogs located in the B chromosomes are transcribed to about similar expression levels as those residing in the A chromosomes, and that the above up-regulations are due to B chromosome transcription. Remarkably, five IC genes (*bmbl, hem1, fas2, rbbp6* and *slnl1*) also showed up-regulation (see Table S3). Although the IC pattern suggests the presence of incomplete paralogs in the B chromosome, our present approach cannot rule out the additional presence of full copies for IC genes, or even a possible functional role through the RNA interference pathway (36).

Sequence analysis of the 25 B-genes revealed high variation between them for the number of nucleotide differences in respect to the A chromosome paralogs. At one extreme, there were 6 genes showing no differences at all, and 6 showing a single difference in the 3’-UTR or the CDS region (see Table 3 and Dataset S2). On the other hand, 6 genes showed 4-31 nucleotide differences in respect to the A paralogs, including 1-3 deletions of 1-14 nucleotides (see Table 3). Finally, the 7 remaining genes showed 2-13 nucleotide substitutions. We interpret these results as reflecting differences in the time of gene arrival to the B chromosome, the oldest genes showing higher differentiation in respect to the A chromosome genes, and the youngest genes being those showing few or no changes at all, in resemblance with the evolutionary strata found on sex chromosome differentiation (37) and the evolution of the germ-line restricted chromosome in songbirds (38). It is plausible to assume that the most important genes for the B chromosome should be those that arrived first, as one or more of them should be crucial for the drive mechanisms that guaranteed the initial invasion of this parasitic element (39). Since B chromosomes in *L. migratoria* show transmission advantage through both sexes, by male mitotic non-disjunction and female meiotic drive (30), the most important genes for a B chromosome like this are those involved in cell cycle regulation. Although we uncovered five of these genes (*apc1, lok, mdm2, rbbp6* and *cdc25*) in the B chromosome, only *apc1* showed signs of being old in the B chromosome as it displays 9 substitutions in respect to the corresponding A chromosome paralog, 8 being synonymous and 1 non-synonymous. The latter change implied the replacement of glutamine (in the Ref sequence) for lysine (in the Alt sequence), and Provean analysis yielded −1.117 score, indicating that this change is putatively neutral for protein function. Interestingly, *mdm2* and *cdc25* lacked nucleotide changes in respect to the A chromosome paralogs, *lok* showed a single synonymous substitution, and *rbbp6* showed a single non-synonymous substitution being apparently neutral after Provean analysis. This suggests the recent arrival of these three genes to the B chromosome (for additional discussion on the age of the B chromosome, see Supplementary Discussion).

The most remarkable gene is thus *apc1*, which codes for the largest subunit of the Anaphase Promoting Complex or Cyclosome (APC/C), a cell cycle E3 ubiquitin ligase regulating the metaphase-anaphase transition during mitosis (40–42), and also playing a role during meiosis (43, 44). During mitosis, APC/C activation is tightly controlled by the spindle assembly checkpoint (SAC) which, in presence of kinetochores being unattached to microtubules, generates the mitotic checkpoint complex (MCC) which inhibits APC/C activation until all chromosomes are properly aligned to the metaphase plate (45, 46). When this condition is met, APC/C is activated and anaphase begins. There is no doubt that the *apc1* gene shows up-regulation in B-carrying individuals, due to the transcription of the gene paralogs residing in the B chromosome. These extra *apc1* transcripts, if translated into proteins, could undermine MCC surveillance for bipolar kinetochore orientation, even if the two B sister kinetochores are monopolarly oriented, thus facilitating the non-disjunction of the B chromosome. In *L. migratoria*, the male B-drive mechanism was first suggested by Nur (25) as a kind of gonotaxis (*sensu* reference 6) by which non-disjunction of the B chromosome during early embryo mitoses where germ line is being determined, leads to B accumulation in germ cells. This fact was later shown by Kayano (26) by comparing B numbers between germ and somatic cells from the same males. Therefore, the key stage where increased amounts of APC/C might play in favor of this kind of B chromosome accumulation, is early embryo development. In fact, the mitotic non-disjunction of this B chromosome was cytologically visualized in three-day-old embryos by Pardo et al. (47). This makes *L. migratoria* a unique model system to study how APC/C amounts influence chromosome segregation, given that the B-derived proteins can be distinguished from the A-derived ones because of the glutamine-lysine substitution.

The second B-drive mechanism in *L. migratoria* takes place during female meiosis, by means of preferential migration of the B chromosome towards the secondary oocyte instead of to the first polar body (30), a mechanism cytologically shown by Hewitt (48) in the grasshopper *Myrmeleotettix maculatus*. In locusts and grasshoppers, meiotic resumption of primary oocytes occurs upon fertilization during egg laying, so that recently laid eggs are at first meiotic metaphase (49). Therefore, the preferential migration of the B chromosome to the secondary oocyte in *L. migratoria* (30) actually occurs while APC/C is operating to promote the metaphase-anaphase transition. It is thus tempting to speculate that the extra amount of APC/C expected in presence of the B chromosome may change checkpoint regulation giving it a chance to perform meiotic drive.

Taken together, our present results show that B chromosome destiny may depend critically on the presence of active genes in it. Darlington and Upcott (4) claimed that “new extra chromosomes appear from time to time in many species, but most of them come to nothing”. We can now complement this sentence by adding that “the remainder can become true B chromosomes if they are genetically well equipped”.

## Material and Methods

(We describe here which procedures were done. For full details on how they were performed, see Supplementary Materials and Methods)

### Materials, nucleic acid isolation and sequencing

We collected 45 males of *Locusta migratoria* at three natural Spanish populations, two in the Cádiz province (Finca El Patrón and Puente de Hierro, at 8 Km distance), and Padul in the Granada province (about 250 Km apart). Testes were cytologically analysed to determine B chromosome presence, and body remains were frozen for DNA (hind leg) and RNA (testis and hind leg) extraction. All three 0B males found (two in Cádiz and one in Padul), along with five B-carrying males (four from Cádiz and one from Padul), were selected for Illumina sequencing. To analyse the possible presence of B chromosome sequences in Asian specimens, we used a gDNA Illumina library previously used for genome assembly by Wang et al. (32), 16 RNA-seq bioprojects of *L. migratoria* obtained from SRA (Table S5). To estimate B chromosome age, we generated an RNA-seq library from one *Oedaleus decorus* male (obtained by us), and also four RNA-seq libraries of *O. asiaticus* obtained from SRA. The ancestry of all these materials was checked by assembling their mitogenomes and building a ML tree along with the sequences used by Ma et al. (35) to build a *L. migratoria* phylogeography (Fig. S4).

### De novo *transcriptome assembly, annotation, mapping and selection*

Bioinformatic procedures to search for B chromosome genes were similar to those described in Navarro-Domínguez et al. (18) and are graphically summarized in Fig. S5. We applied this protocol to the 12 RNA-seq and 6 gDNA libraries from Cádiz males, by generating a *de novo* transcriptome which was used as a reference to map the gDNA reads to detect transcriptome contigs showing overabundance in B-carrying gDNA libraries compared to B-lacking ones, and estimated the number of gene copies per haploid genome. To find contigs being candidate to reside in the B chromosome, we first selected those showing less than four copies per genome (thus discarding highly repetitive elements) and more than 0.5 copies (thus discarding contigs putatively showing coverage problems) (Table S1). Among these contigs, we selected, as possible candidate to be present in the B chromosome, those contigs showing a genomic fold change [gFC= log2(+B/0B)] associated to B chromosome presence, being higher than 0.585 in all four B-carrying individuals analyzed. This threshold was set by assuming that every A or B chromosome carried one copy of the gene.

### Sequence analysis for selected contigs

We first checked if the CDS was complete in each of the selected contigs. If not, we tried to complete it by sequence search in an additional *de novo* transcriptome assembled with the four B-lacking RNA libraries. Using the longest version obtained for all selected contigs, we performed an additional mapping of all genomic and transcriptomic libraries to get estimates of coverage per nucleotide site along the contig. We then calculated the mean, standard deviation (SD) and coefficient of variation (CV) of coverage per nucleotide site in the 0B and +B libraries for each CDS. The CV values were used to generate an index of gene completeness, based on the fold change for variation in coverage per site (cvFC) due to B presence, calculated as log2 of the quotient between +B and 0B CVs. Functional annotation of the selected contigs was performed by Gene Ontology (GO) and Eukaryotic Orthologous Groups of proteins (KOG).

### Transcription analysis of B chromosome genes

We calculated the transcriptomic fold change (tFC) due to B chromosome presence as log2 of the quotient between +B and 0B coverage in the RNA-seq libraries, which provided a first indication of B chromosome gene activity. To reliably identify transcripts coming from B-gene activity, we performed SNP analysis for all selected genes to search for B-specific sequence changes. At each variable nucleotide position, we considered as reference (Ref) the nucleotide being fixed in the 0B libraries, and as alternative (Alt) that being present only in +B gDNA or RNA. To increase the reliability of the SNPs observed, we selected only those variants being present in all four B-carrying gDNA libraries. To go a step further in reliability of our transcription analysis, we defined haplotypes based on SNPs sited a distance lower than 100 nt, i.e. shorter than the 125 nt of the Illumina read size obtained for the gDNA libraries. This allowed defining 2–6 haplotypes, for each of 10 genes, which can be physically found in the Illumina reads. We then scored haplotype frequency (i.e. the proportion of read counts showing it) in each gDNA and RNA library from each male, and calculated per copy gene transcription rate (GTR) of a B-specific haplotype as the quotient between RNA and gDNA read counts.

### Estimation of B chromosome age

We first performed a phylogeographic analysis of the full mitogenomes of all individuals analyzed here, and also those used by Ma et al. (35) for worldwide samples. This showed that the two *L. migratoria* populations from Spain, analyzed here, belong to the southern lineage defined by Ma et al. (35), whereas the *L. migratoria* population from China belong to the northern lineage. We then used BLASTN (34) to search for the sequences of the 10 genes showing B-specific SNPs in the CDS (*ampe, apc1, bmbl, fmo, nrc2a, suox_a, suox_b, tret1, var1* and *zsc22*) in B-carrying and B-lacking libraries from Spain and China, and also in *O. decorus* and *O. asiaticus* which were used as outgroups. We then aligned and concatenated all these sequences and built a phylogram by the Neighbor-Joining method with MEGA v5 software (50), and a Bayesian chronogram with BEAST v1.7 (51), calibrating the node separating the *Locusta* and *Oedaleus* genera in 22.81 mya (52), and the node separating the *L. migratoria* SL and NL in 0.895 mya (35).

### qPCR validation

Genomic overabundance associated with B chromosome presence was tested for 25 genes in five 0B and six +B individuals. Primer pairs anchoring in the same exon were designed with Primer3 (53) preferentially on regions with low sequence variation and high read mapping coverage (Table S9). Quantitative PCR was carried out as described in Navarro-Dominguez et al. (18).

## Supporting information

Main and supplementary tables

## Acknowledgments

The material from Los Barrios (Cádiz) was sampling with the help of Francisco Jiménez-Cazalla and Jorge Doña. We are also grateful to Brian Palestis, Robert Trivers, Neil Jones, Jesper Bolman and Alexander Suh for useful comments to an earlier version of this manuscript. This research was funded by Secretaría de Estado de Investigación, Desarrollo e Innovación (CGL2015-70750-P). FJ Ruiz-Ruano was supported by a Junta de Andalucía fellowship (Spain) and a Lawski scholarship (Sweden).

