## Supplementary Information for "Evolutionary success of a parasitic B chromosome rests on gene content"

**This PDF file includes:**

Supplementary Discussion
Supplementary Materials and Methods
References for SI reference citations
Figs. S1 to S5
Captions for Tables S1 to S9
Captions for Datasets S1 and S2

**Other supplementary materials for this manuscript include the following:**

Tables S1 to S9
Datasets S1 and S2

### Supplementary Discussion

Our estimate of B chromosome age, based on the sequence of ten B chromosome genes, indicates that it arose about 2 mya. Therefore, this B chromosome, which shows drive mechanisms through both sexes (1), has had ample time to reach natural populations in all continents where this species lives (2–8). A previous estimate by Teruel et al. (9), using part of the H3 and H4 histone gene sequences, indicated 750 kya for B age, a figure being close to our lower confidence intervale estimate (880 kya) even though it was based in in only a few hundreds of nucleotide sites. Our present results thus corroborate that B chromosomes in *L. migratoria* are quite old, in resemblance to the 2 my old ones in maize (10) and older than the 1.1-1.3 my old Bs in rye (11).

### Supplementary Materials and Methods

**Materials, nucleic acid isolation and sequencing.** We collected a total of 45 males of *Locusta migratoria* at three natural Spanish populations in Finca El Patrón (Cádiz province) (36.20685N, 5.46481W) (20 males), Puente de Hierro (Cádiz) (36.19251N, -5.55131W) (15 males) and Padul (Granada) (37.01472N, 3.60944W) (10 males). After anesthesia, we fixed one testis in 3:1 ethanol-acetic acid, for cytological analysis, and the other testis and body remains were separately frozen in liquid nitrogen for DNA and RNA extraction. We determined the presence of B chromosomes by squashing two testis follicles in 2% lacto-propionic orcein and visualizing primary spermatocytes at first meiotic prophase or metaphase. Only two males from Cádiz and one from Padul lacked B chromosomes. We selected these three 0B males and six B-carrying males (four from Cádiz and two from Padul) for genomic and transcriptomic analyses. In the six males from Cádiz, we extracted RNA from the testis and one of the hind legs, and DNA from the other hind leg. In the males from Padul, we used testes for cytological analysis and DNA extraction and body remains for RNA extraction. We sequenced gDNA and RNA for the same 0B individual, gDNA for a 2B individual and RNA for a 1B individual. For genomic DNA extraction we used the GenElute Mammalian Genomic DNA Miniprep (Sigma-Aldrich). For RNA extraction, we used the Real Total RNA Spin Plus (Durviz) (in case of hind legs or total body), and the RNeasy Lipid Tissue kit (Qiagen) (in case of testes), following manucfacturer's recommendations.

The six individuals from Cádiz yielded 12 RNA-Seq paired-end libraries by means of the Illumina HiSeq2000 platform (2x101 nt read length) (~8 Gb per library), as well as 6 genomic DNA (gDNA) paired-end libraries by the Illumina HiSeq2500 platform (2x125 nt) (~3x coverage per library). The two males from Padul yielded two gDNA libraries with ~1x coverage and two RNA-Seq libraries with ~7 Gb per library by means of Illumina HiSeq 2000 (2x101 nt). The genomic and transcriptomic libraries from Cádiz were the same previously reported in Ruiz-Ruano et al. (12) (SRA studies SRP079877 and SRP066094), and Padul genomic libraries had previously been reported in Ruiz-Ruano et al. (13), with two additional RNA-Seq libraries (SRA study SRP065963). We used a Illumina HiSeq2000 library (2x100 nt) of gDNA from the China culture used to assembly the genome by Wang et al. (14) with ~44x coverage (SRA library SRR764583).

We also used several RNA-seq libraries to assemble transcriptomes. In 2015, we collected one male of *Oedaleus decorus* (i.e. the *L. migratoria* closest relative living in the Iberian Peninsula) close to Capileira (Granada, Spain) (36.96111N, 3.33583W), extracted RNA from the testes and sequenced it in an Illumina HiSeq2000 platform (2x101 nt) yielding about 9 Gb. In addition, we used the four Illumina HiSeq2000 (2x125 nt) libraries of *O. asiaticus* (~41 Gb in total) included in the SRA study no. SRP059063 (15).

To check the ancestry of the three different *Locusta* and *Oedaleus* projects we assemblied their mitogenomes with MITObim (16) randomly selecting 1 million read pairs with Seqtk (https://github.com/lh3/seqtk) and using, as reference, the sequences with GenBank accession numbers X80245 (*Locusta migratoria*) and EU513374 (*Odealeus asiaticus*) for *Locusta* and *Odealus* reads, respectively. We joined these six sequences to the mitogenomes used for the phylogeography of the species in Ma et al. (17) and aligned then with MAFFT v7 (18) using LINSI options. Then we removed the Control Region since it is incomplete in the RNA-Seq assemblies and built a Maximum Likelihood tree with PHYML v3 (19).

***De novo* transcriptome assembly, annotation, mapping and selection.** Custom scripts written to perform this analysis are deposited in a public repository with instructions to install and launch them (https://github.com/fjruizruano/whatGene).

We used a bioinformatic procedure similar to the protocol described by Navarro-Domínguez et al. (20) to search for genes showing differential abundance between B-carrying and B-lacking libraries (Fig. S5). We generated a *de novo* transcriptome which was used as a reference to map the genomic reads. For this purpose, we concatenated the 12 RNA-Seq libraries and performed an *in silico* normalization with 50x maximum coverage. This selected ~68 million reads which were assembled using Trinity (21) with default options. Since we were only interested in knowing the presence of protein coding genes in the *L. migratoria* B chromosome, we extracted the CDSs being longer than 100 aminoacids, using the Transdecoder software (21) and then reduced redundancy with CDHit-EST (22) with local alignment and the greedy algorithm, and grouped those sequences showing 80% or higher similarity in at least 80% of length (options -M 0 -aS 0.8 -c 0.8 -G 0 -g 1). We annotated the clustered CDSs using RepeatMasker (23) with the RepBase (24) and a custom database of repetitive elements previsously generated by us (12) to annotate contigs for repetitive elements, and the Trinotate pipeline (https://trinotate.github.io) and the SWISS-PROT database (25) to annotate contigs for protein-coding genes.

We then mapped the genomic and transcriptomic reads against the reference transcriptome using SSAHA2 (26). This software allows mapping reads showing high variation in respect to the reference, and it accepts partial read mappings. This is crucial for our purpose, since the sequences used for reference lack introns, which are however present in the genomic reads. We accepted mappings with at least 40 nt with a 80% of minimum identity. We then scored the number of reads mapped per site along the CDS, using the Pysamstats utility (https://github.com/alimanfoo/pysamstats) integrated in the bam_coverage_join.py custom script. This script was also used to calculate the mean, standard deviation (SD) and coefficient of variation (CV) of coverage per site in the 0B and +B libraries for each CDS. In addition, the coverage_graphics.py custom script expressed CDS coverage in each library as the number of copies per haploid genome using the following equation: Copy number= (Coverage x Genome Size)/Library Size. Additionally, we estimated the expression level of the annotated CDSs in the 12 RNA-Seq libraries as RPKM using the following equation: RPKM= ($10^9$*mapped reads)/(total mapped reads*contig legnth).

To normalize genomic coverage, we calculated the genome size of 0B and +B individuals as follows. According to Ruiz-Ruano et al. (27), X chromosome size is 13.63 % the C value and B chromosome size is 18.83% the X chromosome size. In the former paper, a 5.76 Gb C value was calculated for *L. migratoria* as the mean value of several previous estimations in the literature. However, Wang et al. (14) later showed it to be 6.3 Gb after genome sequencing. Therefore, we used this latter C value to recalculate X (0.86 Gb) and B (0.16 Gb) chromosome sizes. considering that a diploid male cell contains two sets of autosomes and a single X chromosome. Therefore, the haploid genome size of a B-lacking male is $G_{0B}$= (2C - X)/2= (2*6.3 - 0.86)/2= 5.87 Gb. We also calculated the average haploid genome size of the four B-carrying individuals from Cádiz ($G_{+B}$) bearing additionally in mind that their mean B frequency was close to 1. Therefore, the haploid genome size for B-carrying individuals is $G_{+B}$= $G_{0B}$ + (B frequency*B size)/2= 5.87 + (1*0.16)/2=

5.95 Gb. We separately estimated mean CDS coverage in both 0B and +B genomic libraries and then we filtered out CDSs according to coverage, excluding highly represented CDSs, because they were candidates to be repeated sequences, by selecting CDSs with a mean copy number lower than 4 in the genomic 0B libraries. In addition, we excluded CDSs with copy number being lower than 0.5, in both genomic 0B and +B libraries, as they might have come from assembly errors. We then calculated the fold change in CDS coverage (gFC), due to B chromosome presence, as log2 of the +B/0B quotient.

To select contigs being candidate to reside in the B chromosome, we used several stepwise criteria. The first and least stringent criterion (applied to +B and 0B averages) implied selecting all those contigs showing gFC≥0.585, assuming the a single-copy gene would show two copies in a 0B genome and three in a genome carrying 1B, also assuming the presence of a single copy in the B chromosome. This would yield an expected genomic 1B/0B ratio of 3/2= 1.5 and thus gFC= $\log_2(1.5)$= 0.585. To decrease the probability of selecting contigs showing variation in coverage between different B-carrying individuals, but still preserve some interesting contigs whose detection might have failed in a single individual due to low coverage, we applied a second filter (applied to individual +B coverages) consisting in selecting contigs with gFC≥0.585 in at least three out of four +B individuals. For this purpose, gFC was calculated by dividing each +B individual coverage between average coverage of the two 0B individuals analyzed. Then we annotated all these contigs by Blast2GO (28) and applied a third filter by selecting those with gFC≥0.585 in all four +B individuals, also preserving those contigs annotated as genes involved in cell cycle functions with gFC≥0.585 in at least three out of the four +B individuals.

**Sequence analysis for selected contigs.** The CDSs meeting the former criteria were submitted to additional analyses. We first checked if the CDS was complete in the contig. If it was incomplete, we used an additional *de novo* transcriptome assembly generated with the four RNA libraries obtained from testes and legs of the two 0B individuals from Cádiz and searched for transcripts being homologous to the selected contigs by BLASTN (29). We performed functional gene annotation using Eukaryotic Orthologous Groups of proteins (KOG), by searching for the predicted protein sequence in the WebMGA server (http://weizhong-lab.ucsd.edu/webMGA/server/kog/).

Using the longest version obtained for all selected CDSs, we performed additional mapping of all genomic and transcriptomic libraries with SSAHA2, to get estimates of coverage per site along the contig. We then averaged these estimates per contig and calculated standard deviation (SD) and coefficient of variation (CV), using the script coverage_graphics.py script, which also provides graphics showing coverage as mean copy number (gDNA) or RPM (RNA) ±1SD. Finally, we calculated a fold change for the variation in coverage per site in +B and 0B libreries (cvFC) as log2 of the quotient between +B and 0B CVs. For genes showing high coverage for part of an exon only, we analyzed the possible tandem-repeated structure by selecting reads showing homology with this region, and clustering and assembling them with RepeatExplorer (30).

**Transcription analysis of B chromosome genes.** As a first estimation of possible up-regulation of some genes due to the presence of some active copies in the B chromosome, we calculated a transcriptomic fold change (tFC) due to B chromosome presence as log2 of the quotient between +B and 0B coverage in the RNA-seq libraries. For sequence-dependent inferences, however, we searched for B-specific sequence changes in the gDNA and RNA libraries. For this purpose, we performed SSAHA2 mappings of gDNA and RNA reads from Cádiz against the sequences of the selected genes, in order to perform a SNP analysis. We first merged the BAM files using SAMtools (31) for the six different conditions, i.e., gDNA, testis RNA and leg RNA, for 0B and +B. We used the custom script snp_calling_bchr.py to search for SNPs variants found at least 2 times in gDNA +B and with zero counts in the gDNA and RNA 0B libraries. At each position, we considered as reference (Ref) the nucleotide being present in the 0B, and as alternative (Alt) that being present only in +B gDNA or RNA. To increase the reliability of the nucleotidic variations observed, we applied an extra filter selecting those variants being present in the four gDNA libraries from Cádiz B-carrying individuals. We also checked if the Alt variant was present in the libraries used by Wang et al. (14) to decipher the genomic DNA sequence of this species. We used this information to generate the A chromosome sequences with the Ref allele and the B chromosome sequences with the Alt allele, using the custom script sequence_ref_alt.py. We then used Geneious to translate them and manually check if they were synonym or non-synonym changes.

Additionally, to perform an still more reliable transcription analysis, we defined haplotypes based on SNPs sited at distance lower than 100 nt, i.e. shorter than the 125 nt of the Illumina read size obtained for Cádiz gDNA libraries. This yielded a collection of haplotypes based on the physical position of SNPs pairs within a same read. We selected these regions and extracted the reads carrying these haplotypes from the BAM files with the custom script extract_seq_regions.py. Then we aligned the reads with Geneious and scored the number of occurrences for Ref and Alt haplotypes. This allowed defining 2-6 haplotypes, for each of 10 genes, which can be physically found in the Illumina reads, thus avoiding to score possible sequencing errors. By means of read counts in gDNA and RNA of B-carrying individuals it is possible to estimate haplotype frequency (i.e. the proportion of read counts showing it) in gDNA and RNA from the same individuals, and calculated per copy gene transcription rate (GTR) of a B-specific haplotype as the quotient between RNA and gDNA read counts.

**Estimation of B chromosome age.** Prior to the estimation of B chromosome age, we checked the phylogenetic relationship between the different samples of *Locusta migratoria* from China (14), *Oedaleus asiaticus* (15) available in the SRA database, as well as our *O. decorus* sequencing (see above for details), in addition to our *L. migratoria* samples. For this, we generated a consensus mitogenome per bioproject within each species (Fig. S4). This showed that the two *L. migratoria* populations from Spain, analyzed by us, belong to the southern lineage defined by Ma et al. (17), whereas the *L. migratoria* population from China belong to the northern lineage. Once determined the phylogenetic relationships between samples, we used BLASTN (29) to search in the assemblies for the sequences of the 10 genes found in the B chromosome which also showed high coverage along the whole transcript and displaying SNPs in the CDS: *ampe*, *apc1*, *bmbl*, *fmo*, *nrc2a*, *suox_a*, *suox_b*, *tret1*, *var1* and *zsc22*. Then we assembled the transcriptomic reads of *O. decorus* and *O. asiaticus*, applying the same protocol described above for the *L. migratoria* Spanish individuals, and used BLASTN (29) to search for the sequences of the 10 genes in the assemblies. Since we found B-specific variation in the female sequenced by Wang et al. (14) for the genome assembly, we used this mappings against the selected genes to get the Ref and Alt sequences from the northern lineage. This female came from a highly inbred line, so we assumed that intraindividual SNP variation is due to B chromosome presence. We used the mappings from the two 0B individuals from Cádiz to identify the Ref and Alt nucleotides in variable positions, considering only SNPs with at least an Alt/Ref proportion of 10%. Additionally, all positions being fixed in the China individual for a nucleotide being different from that in the Cádiz population, were considered for both Alt and Ref sequences in China.

We then aligned these four sequences with those observed in A and B chromosomes of the *L. migratoria* SL, previously obtained gene by gene, then extracted the CDS and finally concatenated all genes with Geneious v4.8 (32). We built a phylogram by the Neighbor-Joining method with MEGA v5 software (33), applying pairwise deletion, the Kimura 2-parameters substitution model and a bootstrap of 1000 replicates. We built a bayesian chronogram with BEAST v1.7 (34) calibrating the node separating the *Locusta* and *Oedaleus* genera in 22.81 mya (35), and the node separating the *L. migratoria* SL and NL in 0.895 mya (17). We launched three MCMC during $10^8$ generations sampling a tree each 1000 generations. We checked for the convergence of these three runs with Tracer and combined the resulting trees with LogCombiner applying a 10% burnin.

**qPCR validation.** The genomic overabundance associated with B chromosome presence was tested for 25 genes in five 0B and six +B individuals, including the Cadiz individuals with gDNA sequenced by Illumina.

Primer pairs anchoring in the same exon were designed with Primer3 (36). We search for exon limits using the Exonerate software (37) and aligning the transcript sequence against the *L. migratoria* genome assembled by Wang et al. (14). We preferably selected regions with low sequence variability and high read mapping coverage. Primer sequences and amplicon length are shown in Table S7.

Quantitative PCR was carried out as described in Navarro-Dominguez et al. (20). qPCRs were performed on a Chromo 4 Real Time PCR thermocycler (Biorad). Each reaction mixture contained 25 ng of gDNA in 5 µl, 5 µl of SensiMix SYBR Kit (Bioline) and 2.5 µl of each 2.5 µM primer. Reactions were carried out in duplicate. We estimated the amplification efficiency (E) of each primer pair by means of a standard curve performed on a 10-fold dilution series of *L. migratoria* gDNA mixture from the 11 quantified individuals. This gDNA pool was also used as an external calibrator for the qPCR reactions. The relative abundance of each gene in each sample was calculated according to $RQ = E^{CtC-CtS}$ , where RQ = Relative quantity, E = Amplification efficiency (fold increase per cycle), CtC = Ct value of the calibrator sample and CtS = Ct value of each sample.

The Shapiro-Wilks test was used to verify the normality of the RQ values ($p>0.2$), and consequently we applied a Student's t-test to assess statistical significance of differences between the 0B and +B samples.

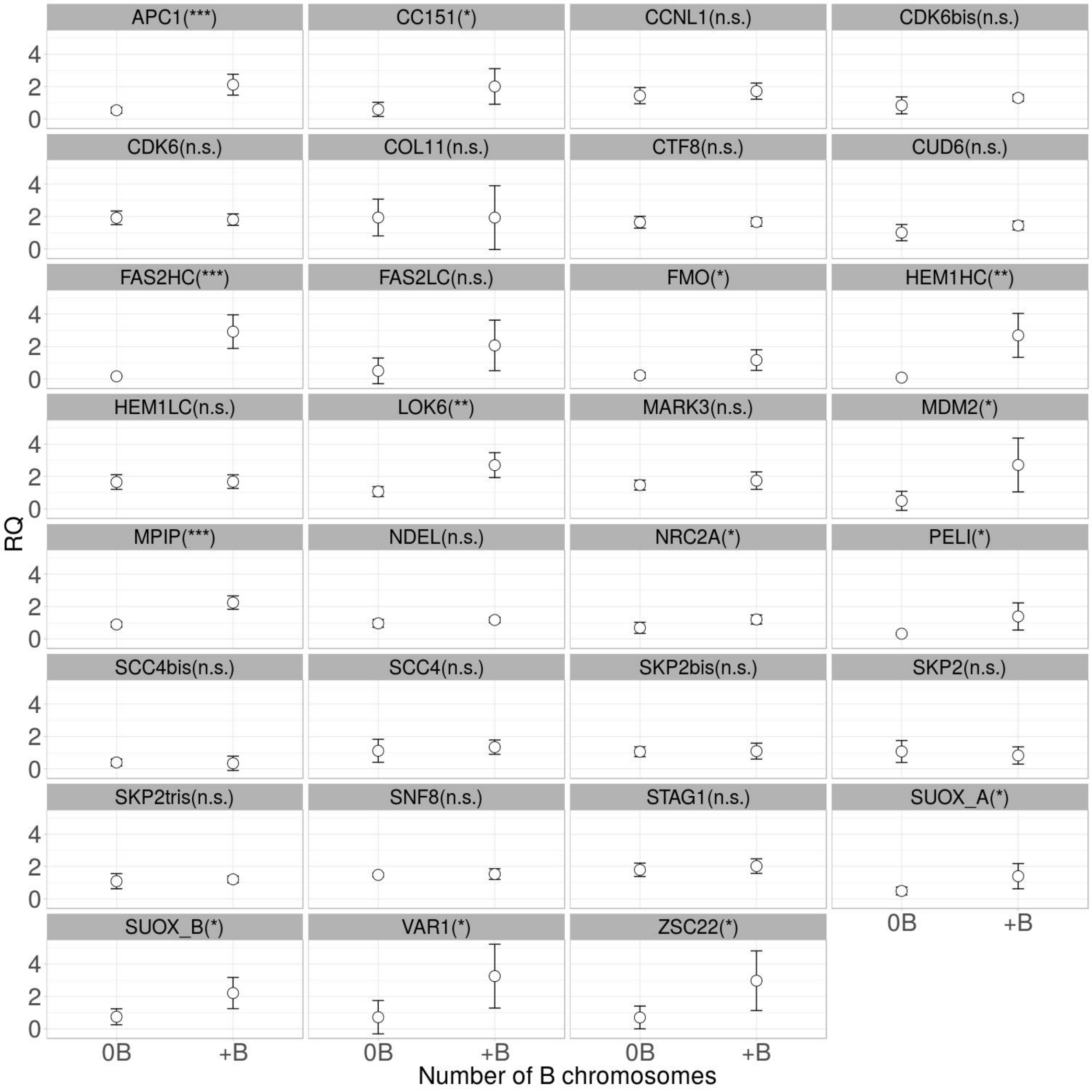


**Fig. S1**. Results for all the qPCR analyses performed on genomic DNA. ***: p<0.001. **: p<0.01. *: p<0.05. n.s.: p>0.05.

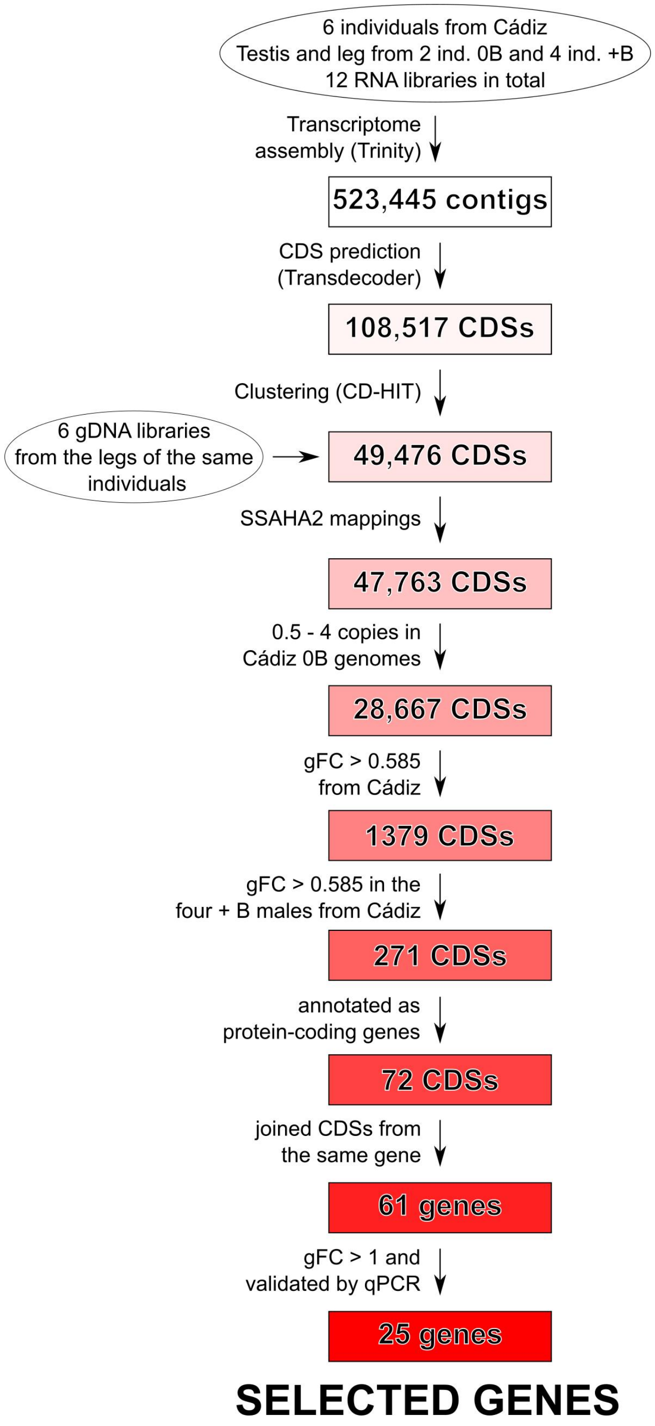


**Fig. S2**. Pathway for gene search in the *L. migratoria* B chromosome.

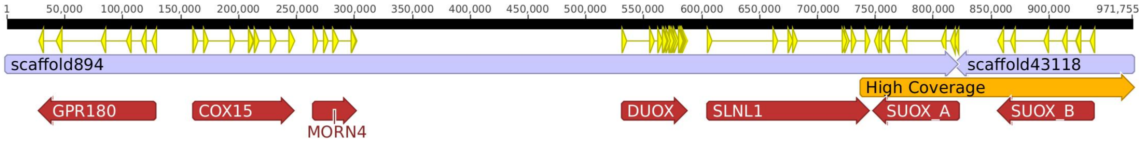


**Fig. S3**. Two scaffolds from the *L. migratoria* genome published by Wang et al. (14) shared the *suox_a* gene, which allowed us to infer that *slnl1*, *suox_a* and *suox_b* are physically linked in the genome. Highlighted in orange color is the region found in the B chromosome, including *suox_a*, *suox_b* and the last exon of the *slnl1* gene.

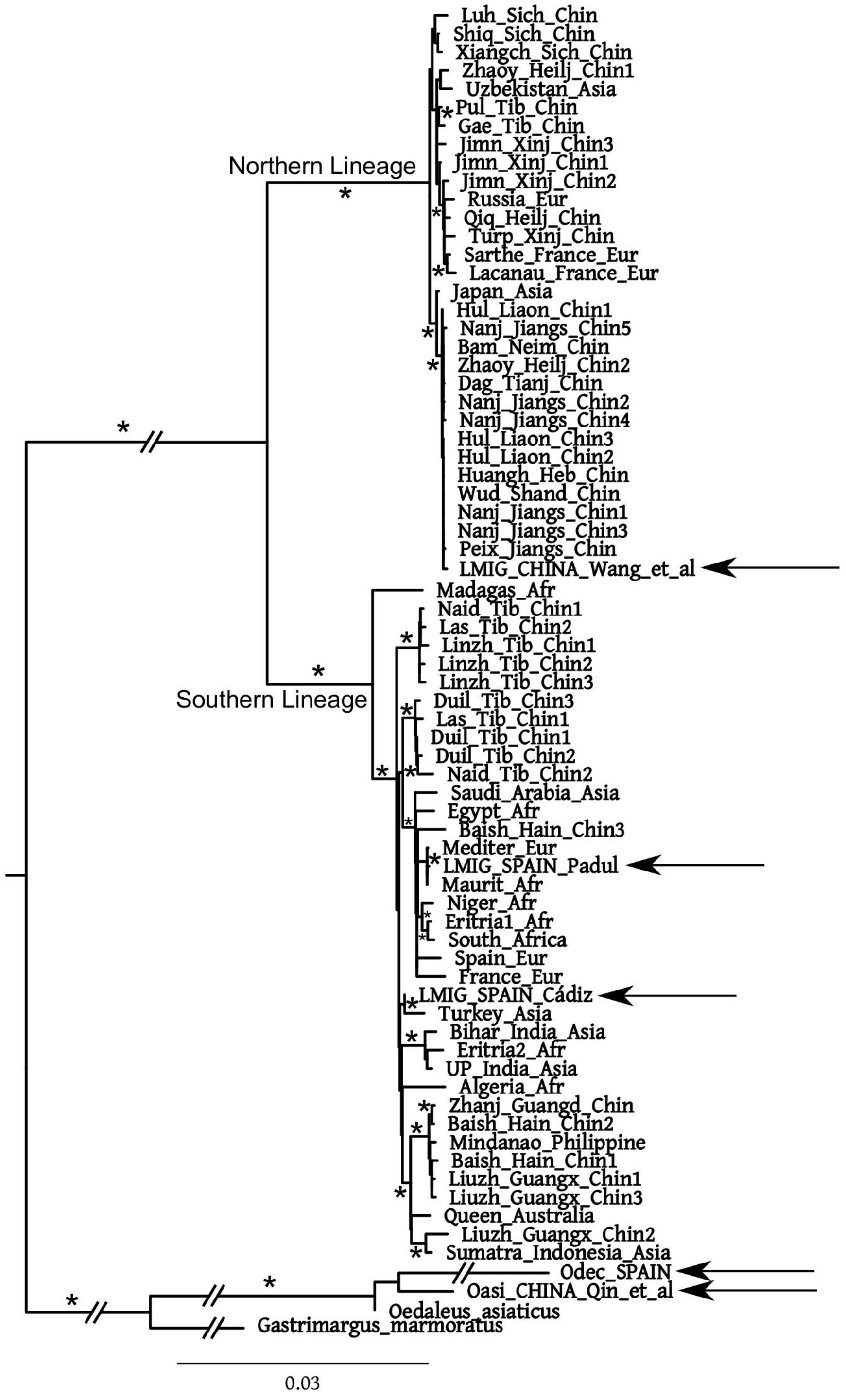


**Fig. S4**. PhyML tree of *Locusta* and *Oedaleus* mitogenomes. Bootstraps equal or higher than 90% are indicated with an asterisk. Arrows point the six mitogenomes assembled for the present work whereas the remainder were those analyzed in Ma et al. (17). Note that the two Chinese populations belong to the *L. migratoria* northern lineage (NL) and the two Spanish populations belong to the southern lineage (SL).

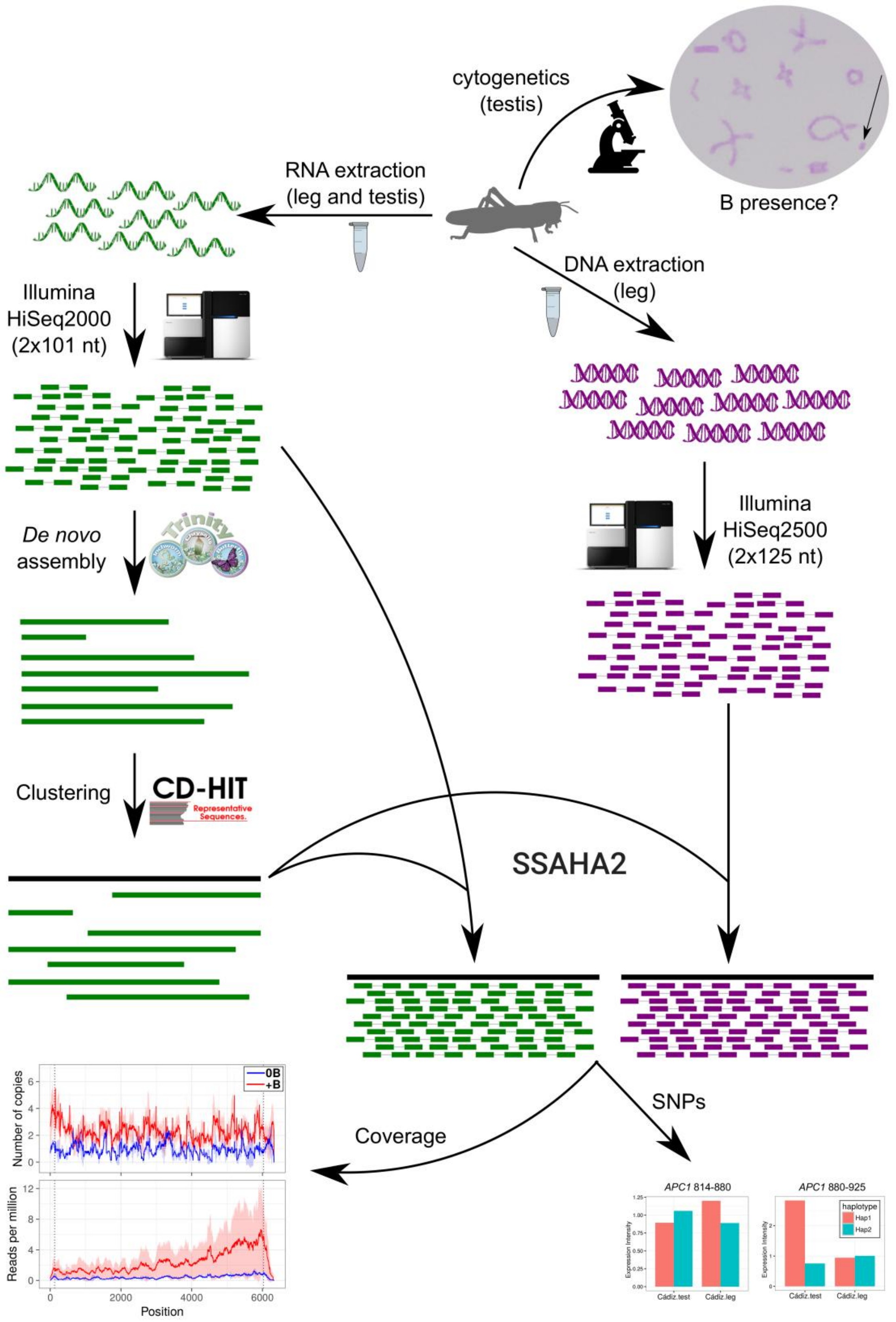


**Fig. S5.** Bioinformatic protocol applied for gene selection, and coverage and SNP analyses. See Supplementary Materials and Methods for details.

### Caption for Tables S1 to S9

**Table S1.** Contigs selected for thorough sequence analysis because they showed gFC≥0.585 in all four +B individuals..

**Table S2.** List of genes sorted per decreasing gFC value in the high coverage region (hc_gFC), and qPCR results for 25 genes showing from 0.06 to 6.14 hc_FC values. Mean coverages were calculated as the average coverage of all CDS nucleotides, and the coefficient of variation (CV) of nucleotidic coverage in +B and 0B were used to calculate a coverage variation fold change (cvFC) as log2(cv+B/cv0B). Negative values of this parameter indicate uniform coverage (UC) and suggest the possibility that the gene CDS is complete in the B chromosome, whereas positive values indicate irregular coverage and suggest the existence of pseudogenical copies in the B chromosome. HC= High coverage, LC= Low coverage. In some genes, two or three different primer pairs were tested by qPCR.

**Table S3.** Genomic (gFC) and transcriptomic (tFC) fold change [log2(+B/0B)] for B chromosome genes. SNPs analysis showing Presence of deleterious aminoacid changes (PDAC) and expression intensity (EI) for SNP-haplotypes. GC= gDNA from Cádiz. GP= gDNA from Padul. Gch= gDNA from China. RTC= RNA-Seq on testis from Cádiz. RLC= RNA-Seq on leg from Cádiz. RP= RNA-Seq from Padul.

**Table S4.** Counts of reference (Ref) and alternative (Alt= B-specific) variants in all gDNA and RNA libraries analyzed in Cádiz.

**Table S5.** RNA-Seq libraries from *L. migratoria* available in SRA database used for SNP calling. Index is the letter used in Table S6 for each bioproject.

**Table S6.** Counts of Ref and Alt haplotypes in all the studied group of libraries, including SRA transcriptomes, for the 169 SNPs specific of +B libraries of Cádiz gDNA and RNA.

**Table S7.** Transcription intensity (TI) of B-specific haplotypes defined by SNPs being at distances lower than one Illumina read (i.e. 100 nt). "0B_C"= B-lacking males from Cádiz; "+B_C"= B-carrying males from Cádiz; TES= Testis, LEG= Hind leg. Note the presence of most B-specific haplotypes in the genome analyzed in China by Wang et al. (2014). Hap 1 always refer to the haplotype present in the A chromosomes..

**Table S8.** Summary of Ref and Alt haplotypes found in +B individuals from Cádiz

**Table S9.** Primer pairs used for qPCR validation.

### Caption for Datasets S1 and S2

**Dataset S1.** Spreadsheet showing the procedure for CDS selection by applying four consecutive filters: i) Among the 47,763CDSs showing mappings for the Cádiz gDNA libraries, we selected those with average number of copies in the 0B libraries between 0.5 and 4 (28,667 CDSs), ii) we then selected those CDSs showing genomic fold change due to B chromosome presence [gFC= log2(+B/0B)] higher than 0.585 (1,379 CDSs), iii) CDS showing gFC> 0.585 in all four +B individuals (271 CDSs) were selected, and iv) 72 CDSs out of 271 were annottated for protein-coding genes.

**Dataset S2.** Coverage graphical display for the 25 protein-coding genes found in the *L. migratoria* B chromosomes from the Spanish individuals collected at Cádiz (Ca) and Padul (Pa), both belonging to the southern lineage (SL), most of which show evidence to be also present in Chinese individuals belonging to the northern lineage (NL). gDNA coverage is shown as number of copies, and RNA coverage is shown as reads per million of mapped reads. Additionally, we add a track showing the position of the B-specific SNPs found in the Cádiz libraries. The coverage patterns inferred from the fold change in coverage variation along CDS length [cvFC= log2(cv+B/cv0B)] are also specified. Note that negative cvFC values indicate uniform coverage (UC) whereas positive values suggest irregular coverage (IC) patterns. The latter pattern could be due to regional amplification (IC-ra), truncation (IC-tr) or satellite DNA formation (IC-sat).
