## Supplementary material for "Evolutionary success of a parasitic B chromosome rests on gene content": Dataset S2

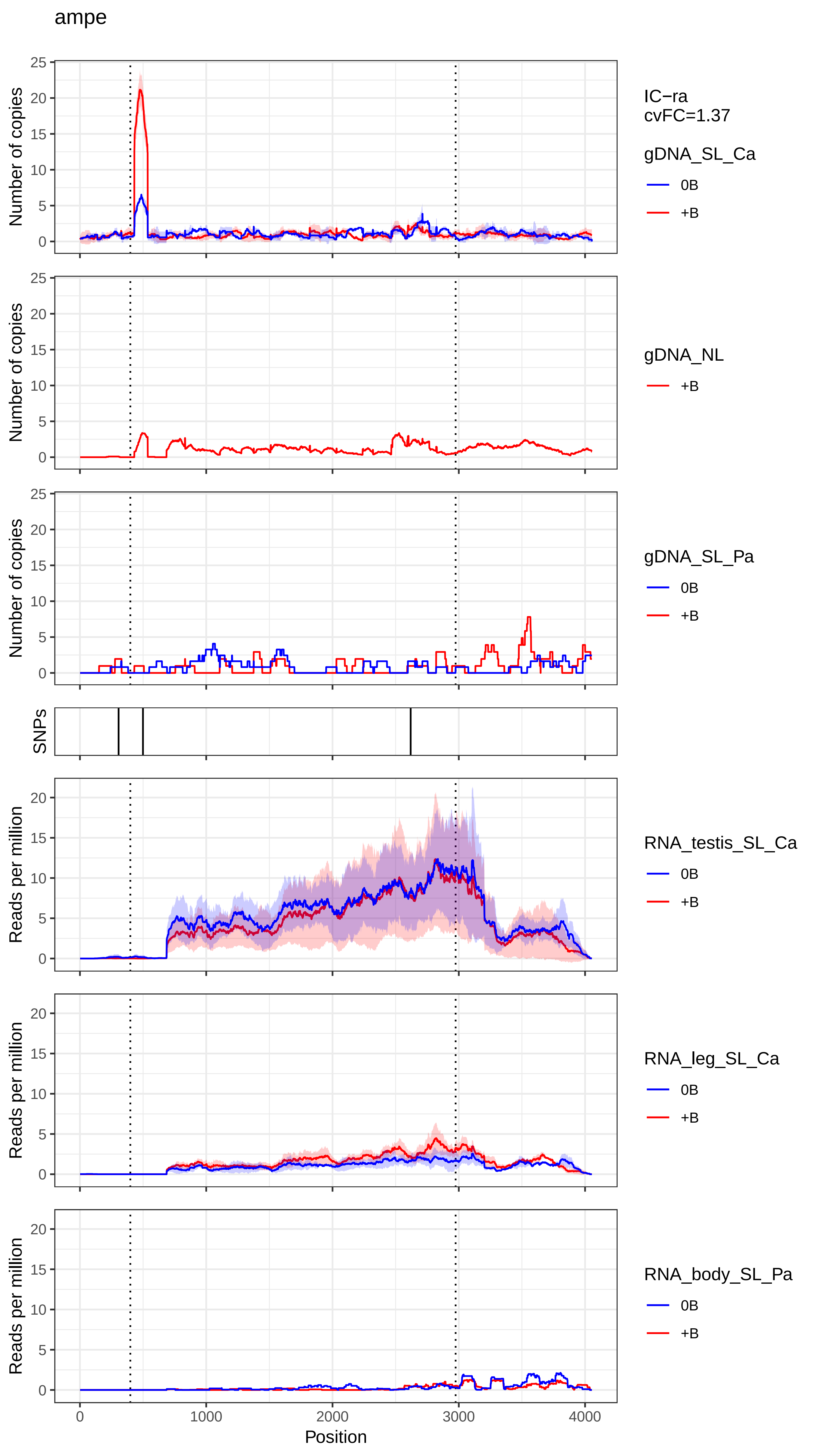

ampe
IC−ra
cvFC=1.37
gDNA_SL_Ca
0B
+B
gDNA_NL
+B
gDNA_SL_Pa
0B
+B
RNA_testis_SL_Ca
0B
+B
RNA_leg_SL_Ca
0B
+B
RNA_body_SL_Pa
0B
+B
Number of copies
SNPs
Reads per million
Position
0
1000
2000
3000
4000

apc1

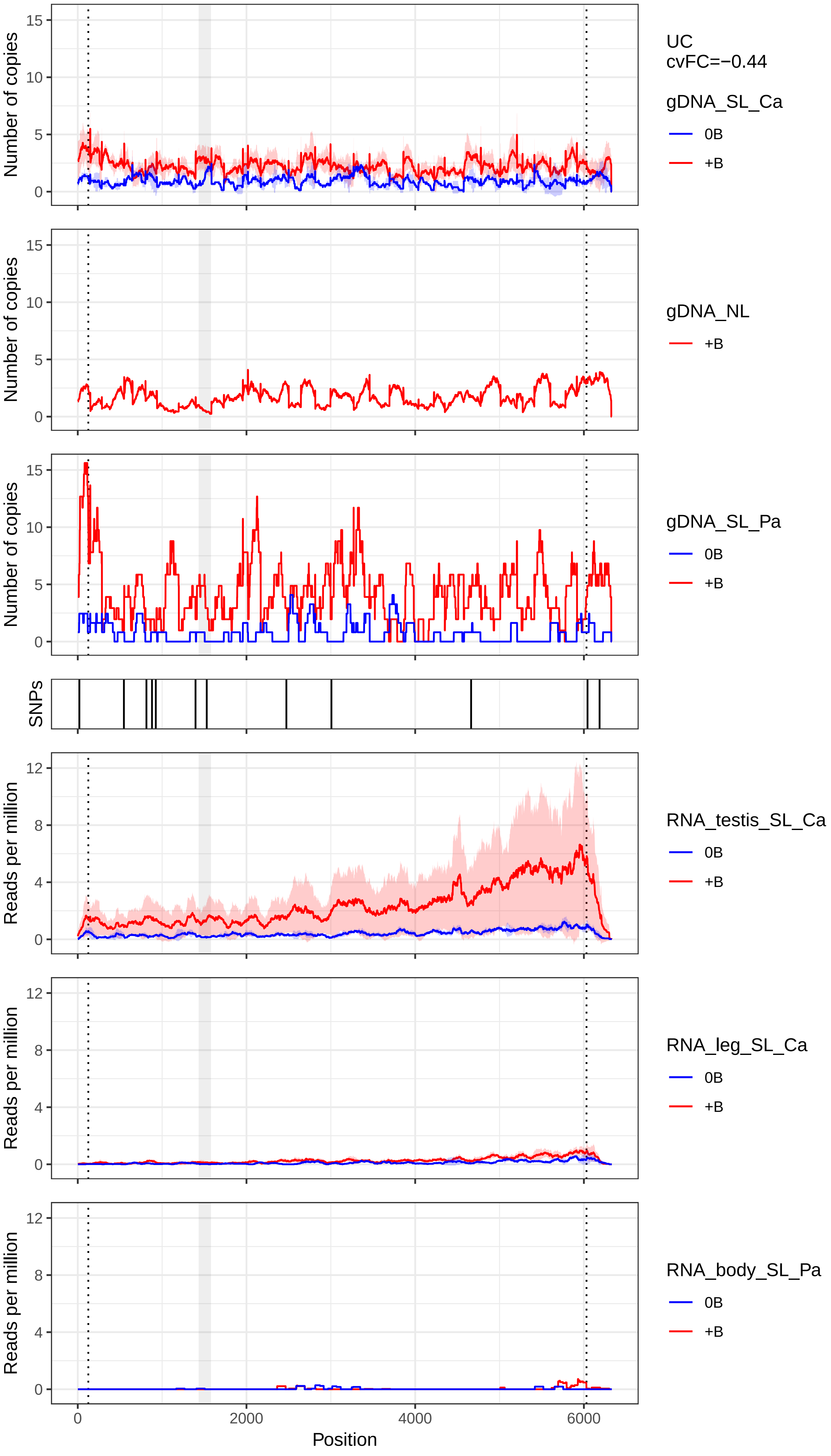

Number of copies
Reads per million
SNPs
Position
UC
cvFC=−0.44
gDNA_SL_Ca
gDNA_NL
gDNA_SL_Pa
RNA_testis_SL_Ca
RNA_leg_SL_Ca
RNA_body_SL_Pa
0B
+B

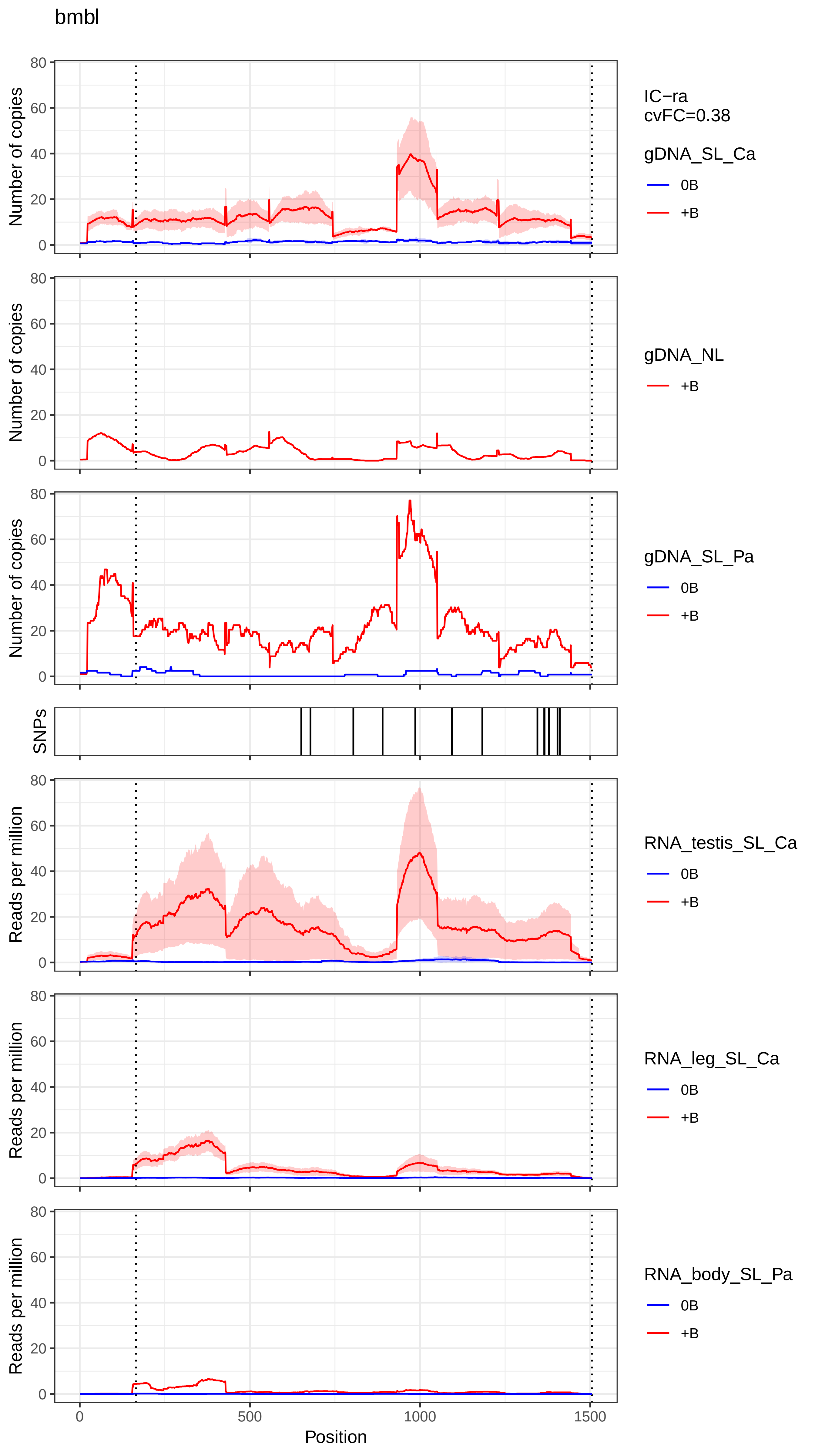

bmbl
IC−ra
cvFC=0.38
gDNA_SL_Ca
0B
+B
gDNA_NL
+B
gDNA_SL_Pa
0B
+B
SNPs
RNA_testis_SL_Ca
0B
+B
RNA_leg_SL_Ca
0B
+B
RNA_body_SL_Pa
0B
+B
Number of copies
Reads per million
Position
0
500
1000
1500

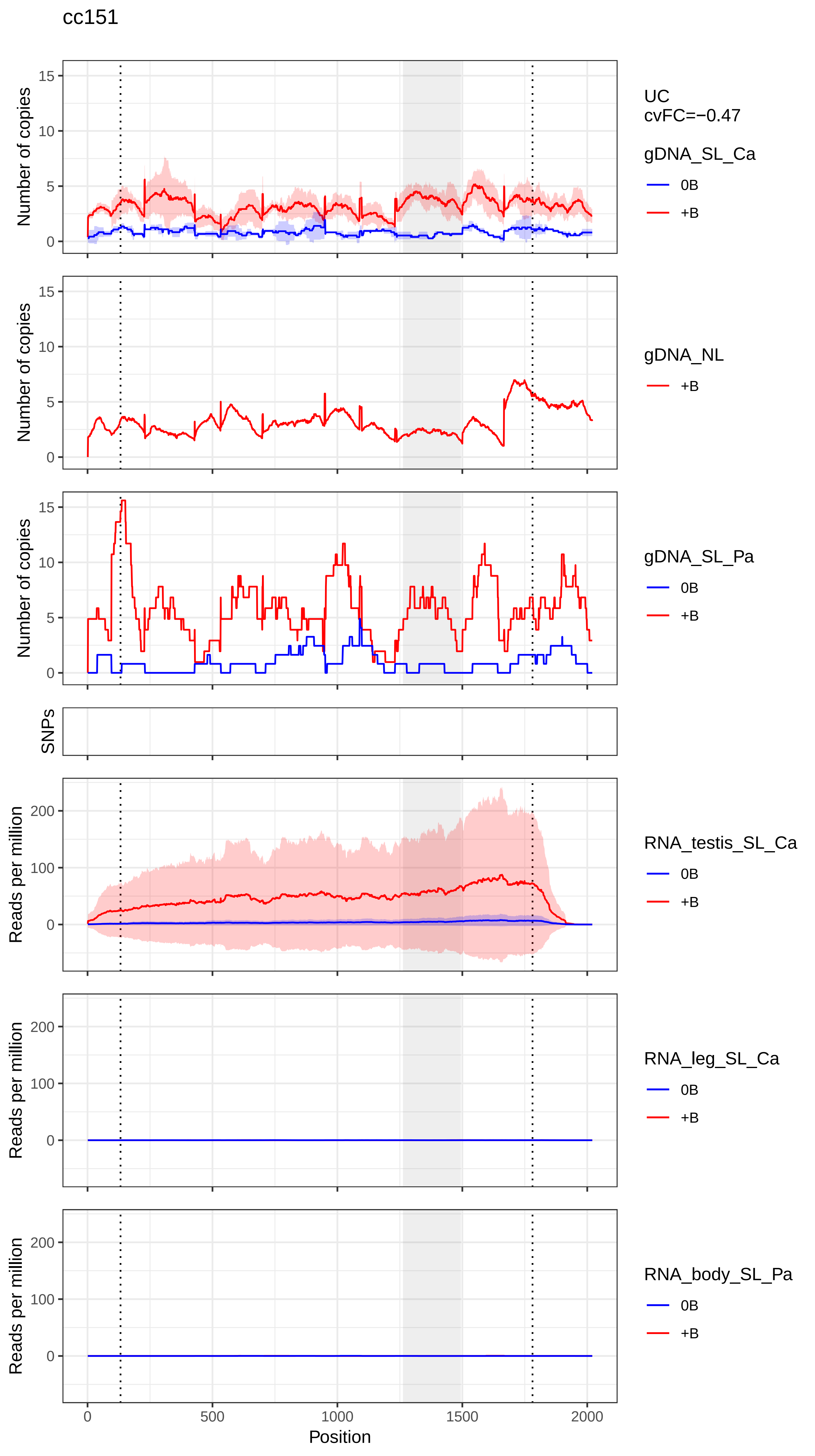
cc151
UC
cvFC=−0.47
gDNA_SL_Ca
0B
+B
gDNA_NL
+B
gDNA_SL_Pa
0B
+B
RNA_testis_SL_Ca
0B
+B
RNA_leg_SL_Ca
0B
+B
RNA_body_SL_Pa
0B
+B
Number of copies
SNPs
Reads per million
Position
0
500
1000
1500
2000
0
5
10
15
100
200

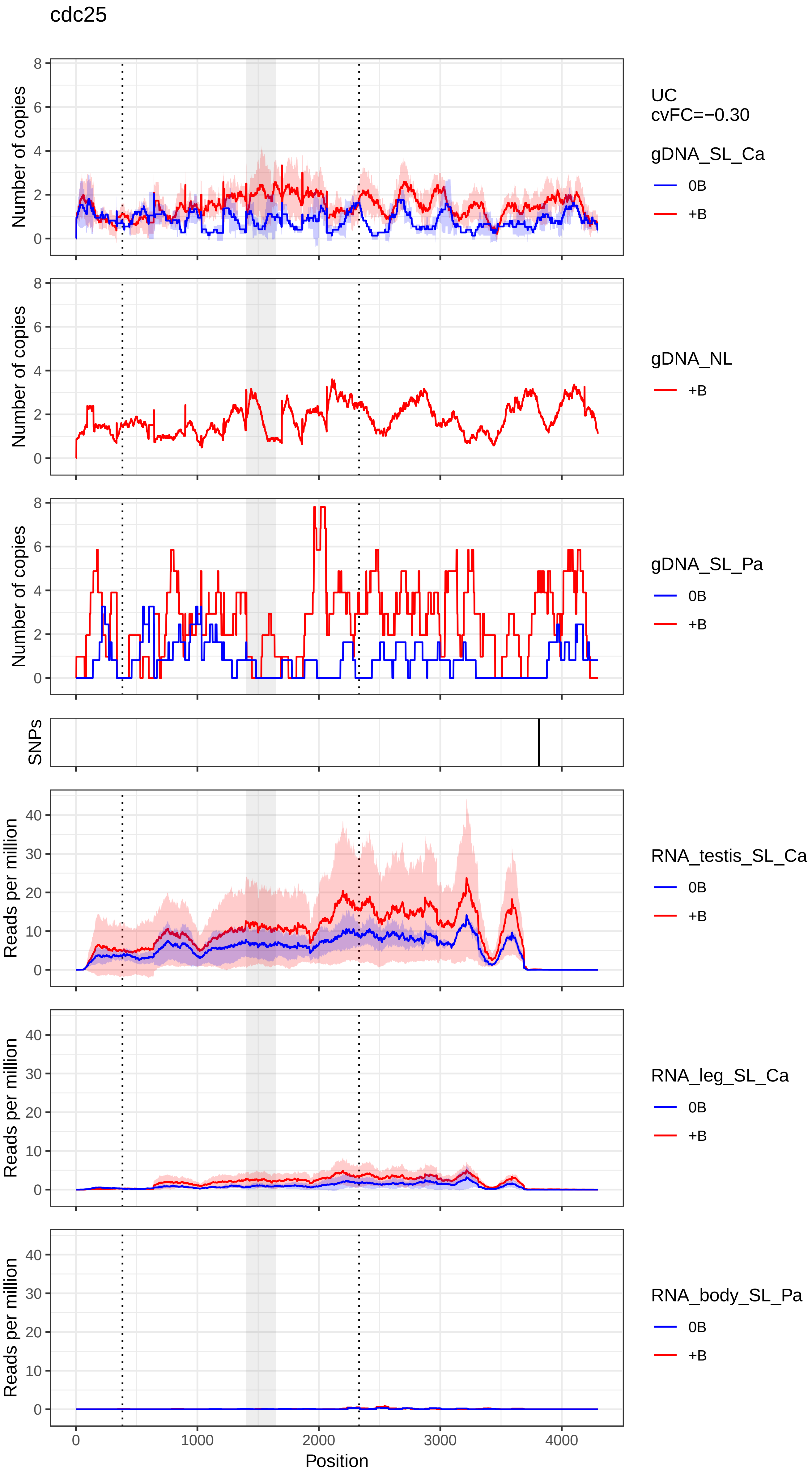
cdc25
UC
cvFC=−0.30
gDNA_SL_Ca
0B
+B
gDNA_NL
+B
gDNA_SL_Pa
0B
+B
SNPs
RNA_testis_SL_Ca
0B
+B
RNA_leg_SL_Ca
0B
+B
RNA_body_SL_Pa
0B
+B
Number of copies
Reads per million
Position
0
1000
2000
3000
4000

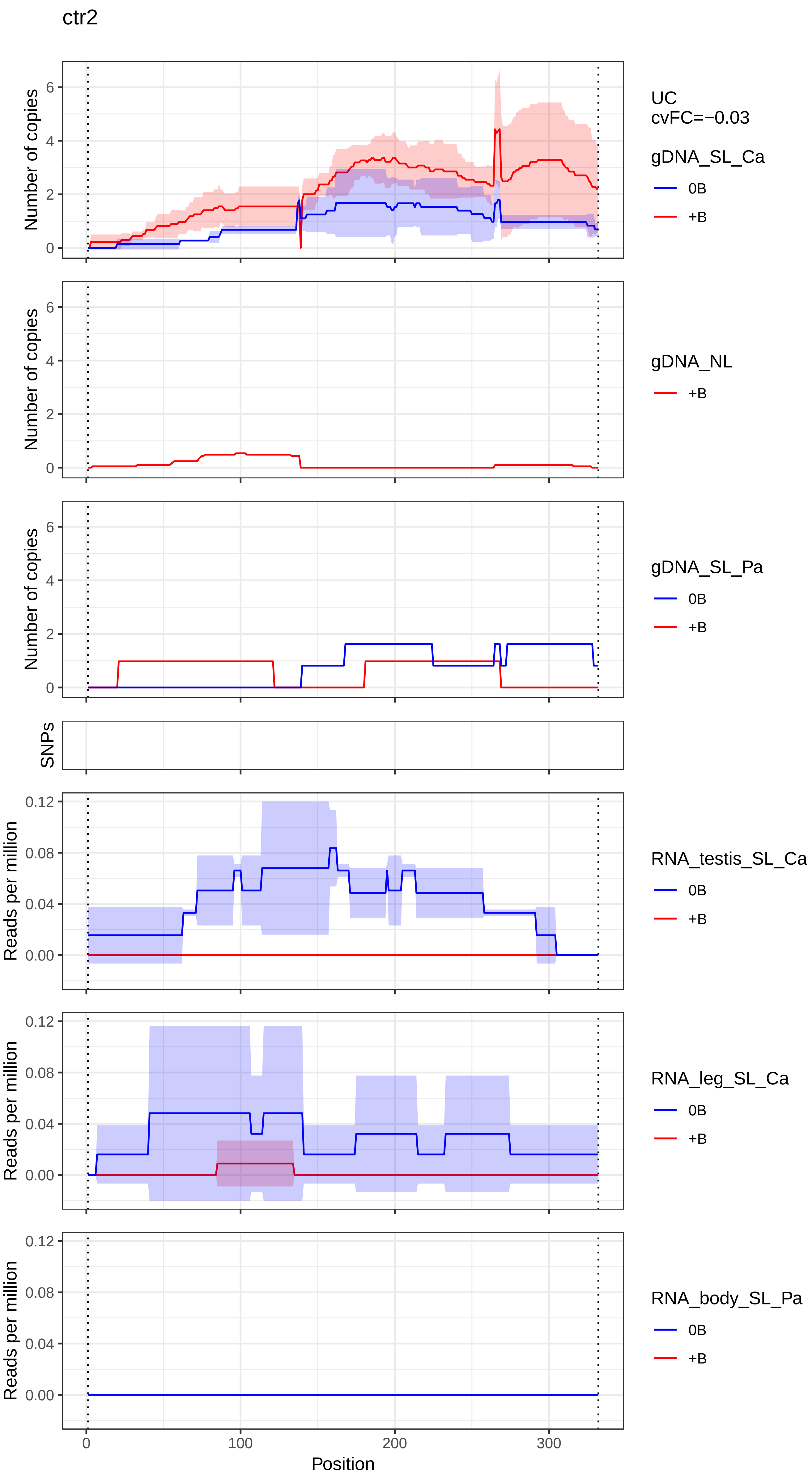
ctr2
Number of copies
UC
cvFC=−0.03
gDNA_SL_Ca
0B
+B
gDNA_NL
+B
gDNA_SL_Pa
0B
+B
SNPs
Reads per million
RNA_testis_SL_Ca
0B
+B
RNA_leg_SL_Ca
0B
+B
RNA_body_SL_Pa
0B
+B
Position

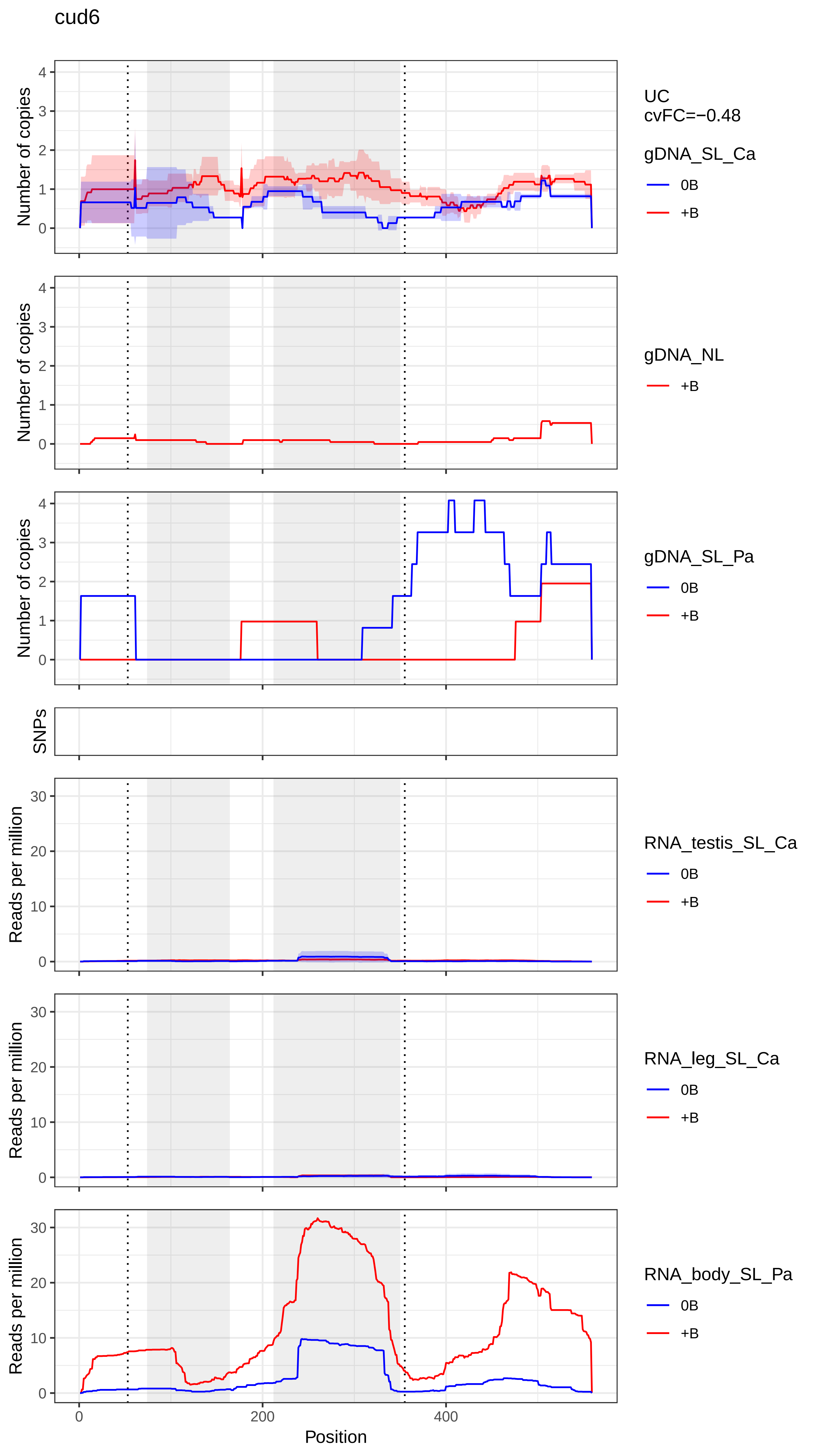
cud6
UC
cvFC=−0.48
gDNA_SL_Ca
0B
+B
gDNA_NL
+B
gDNA_SL_Pa
0B
+B
SNPs
RNA_testis_SL_Ca
0B
+B
RNA_leg_SL_Ca
0B
+B
RNA_body_SL_Pa
0B
+B
Number of copies
Reads per million
Position
0
200
400
0
1
2
3
4
0
10
20
30

fas2

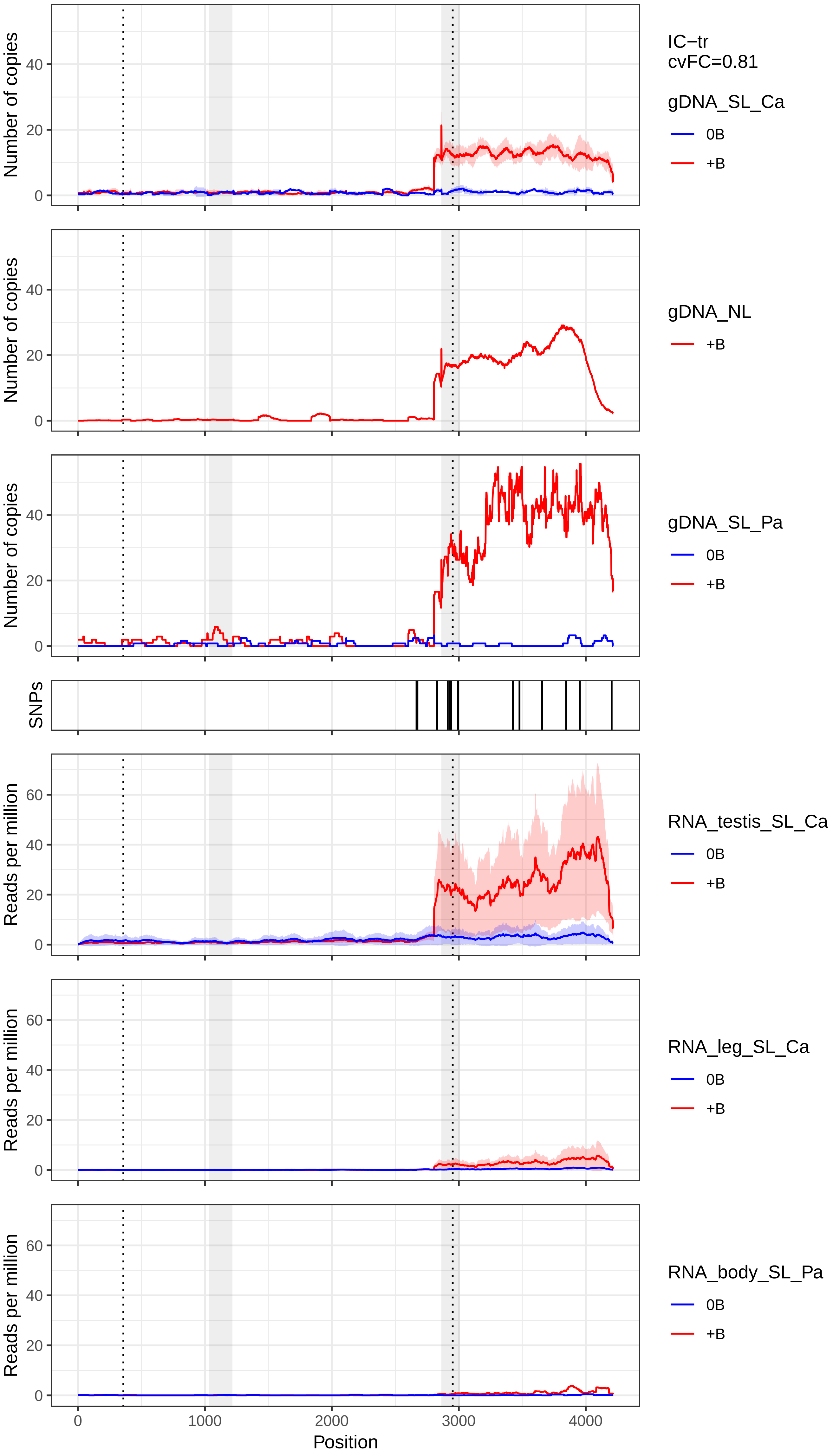

IC−tr
cvFC=0.81
gDNA_SL_Ca
0B
+B
gDNA_NL
+B
gDNA_SL_Pa
0B
+B
SNPs
RNA_testis_SL_Ca
0B
+B
RNA_leg_SL_Ca
0B
+B
RNA_body_SL_Pa
0B
+B
Number of copies
Reads per million
0
20
40
60
0
1000
2000
3000
4000
Position

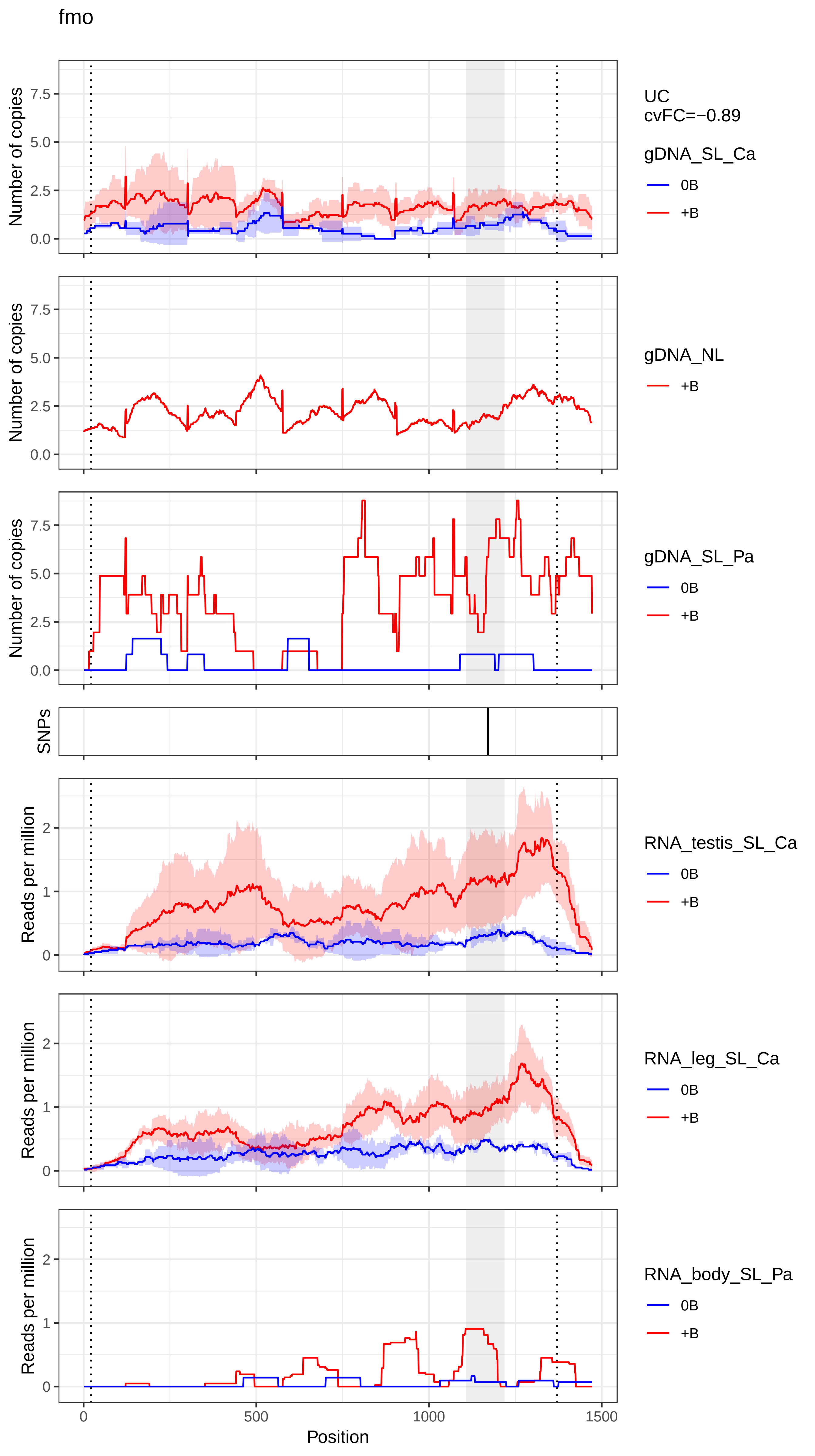
fmo
UC
cvFC=−0.89
gDNA_SL_Ca
0B
+B
gDNA_NL
+B
gDNA_SL_Pa
0B
+B
SNPs
RNA_testis_SL_Ca
0B
+B
RNA_leg_SL_Ca
0B
+B
RNA_body_SL_Pa
0B
+B
Number of copies
Reads per million
0.0
2.5
5.0
7.5
0
1
2
0
500
1000
1500
Position

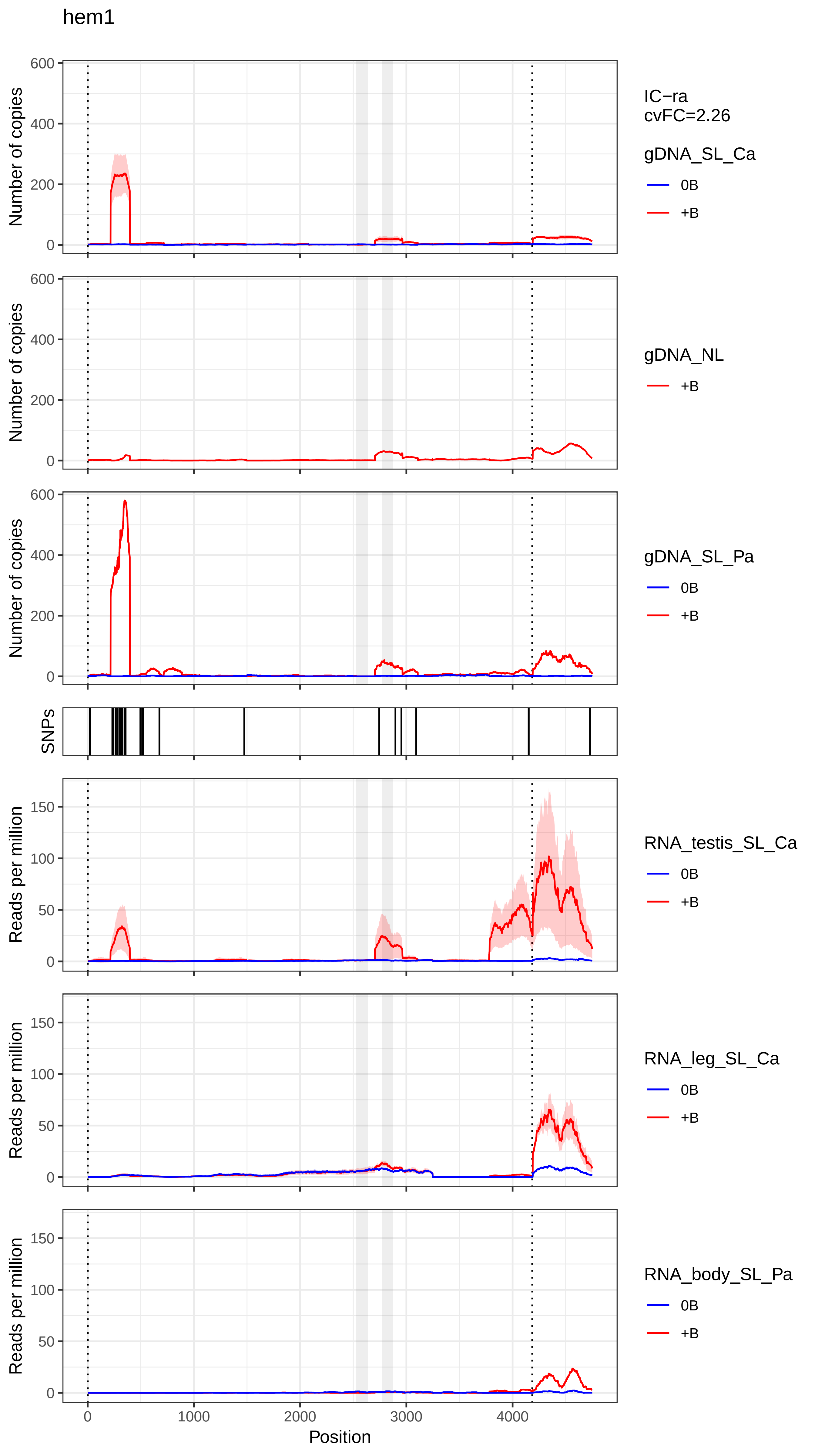
hem1
IC−ra
cvFC=2.26
gDNA_SL_Ca
0B
+B
gDNA_NL
+B
gDNA_SL_Pa
0B
+B
SNPs
RNA_testis_SL_Ca
0B
+B
RNA_leg_SL_Ca
0B
+B
RNA_body_SL_Pa
0B
+B
Number of copies
Reads per million
Position
0
1000
2000
3000
4000
600
400
200
150
100
50

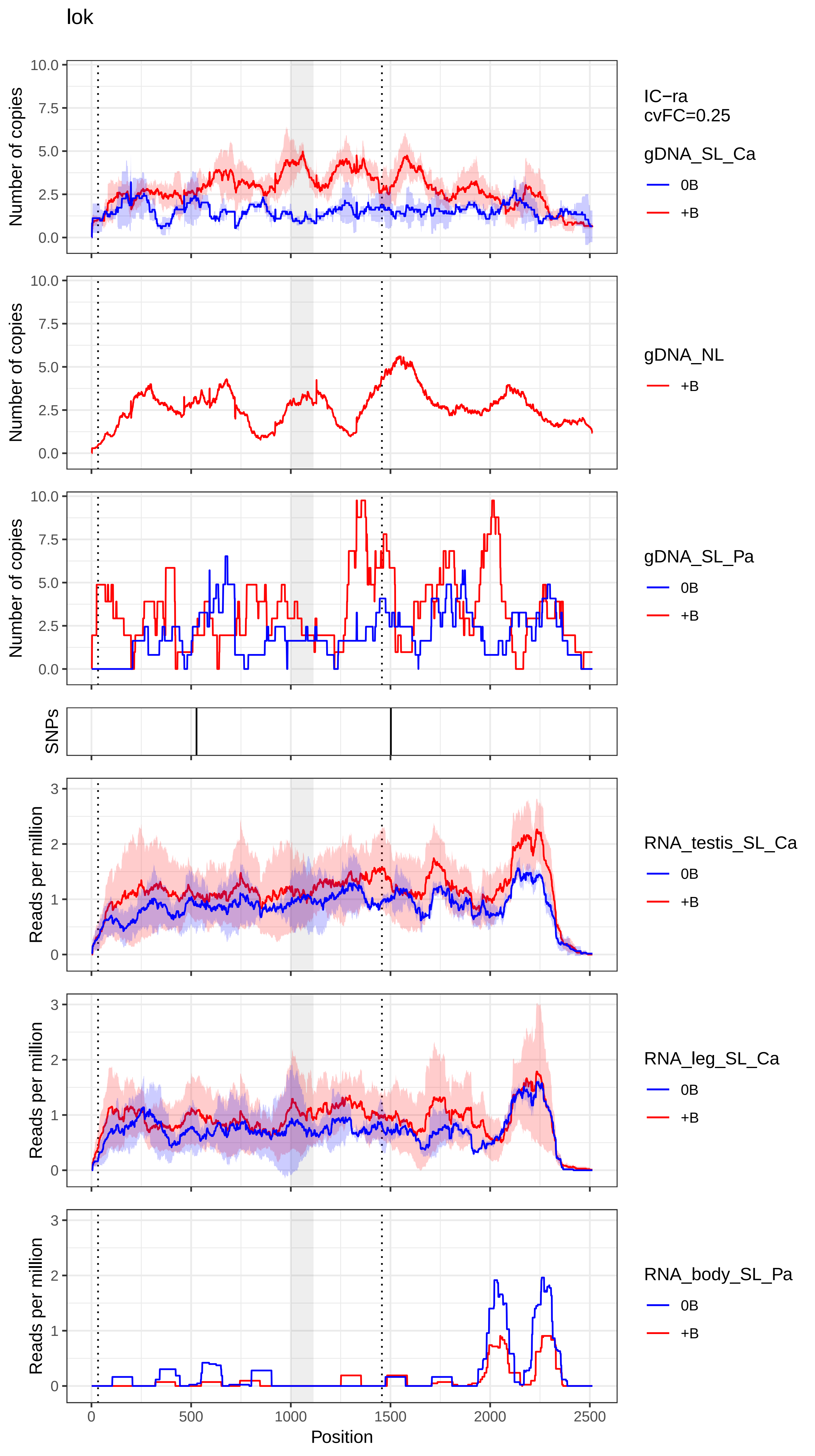
lok
IC−ra
cvFC=0.25
gDNA_SL_Ca
0B
+B
gDNA_NL
+B
gDNA_SL_Pa
0B
+B
RNA_testis_SL_Ca
0B
+B
RNA_leg_SL_Ca
0B
+B
RNA_body_SL_Pa
0B
+B
Number of copies
SNPs
Reads per million
Position
0
500
1000
1500
2000
2500
0.0
2.5
5.0
7.5
10.0
0
1
2
3

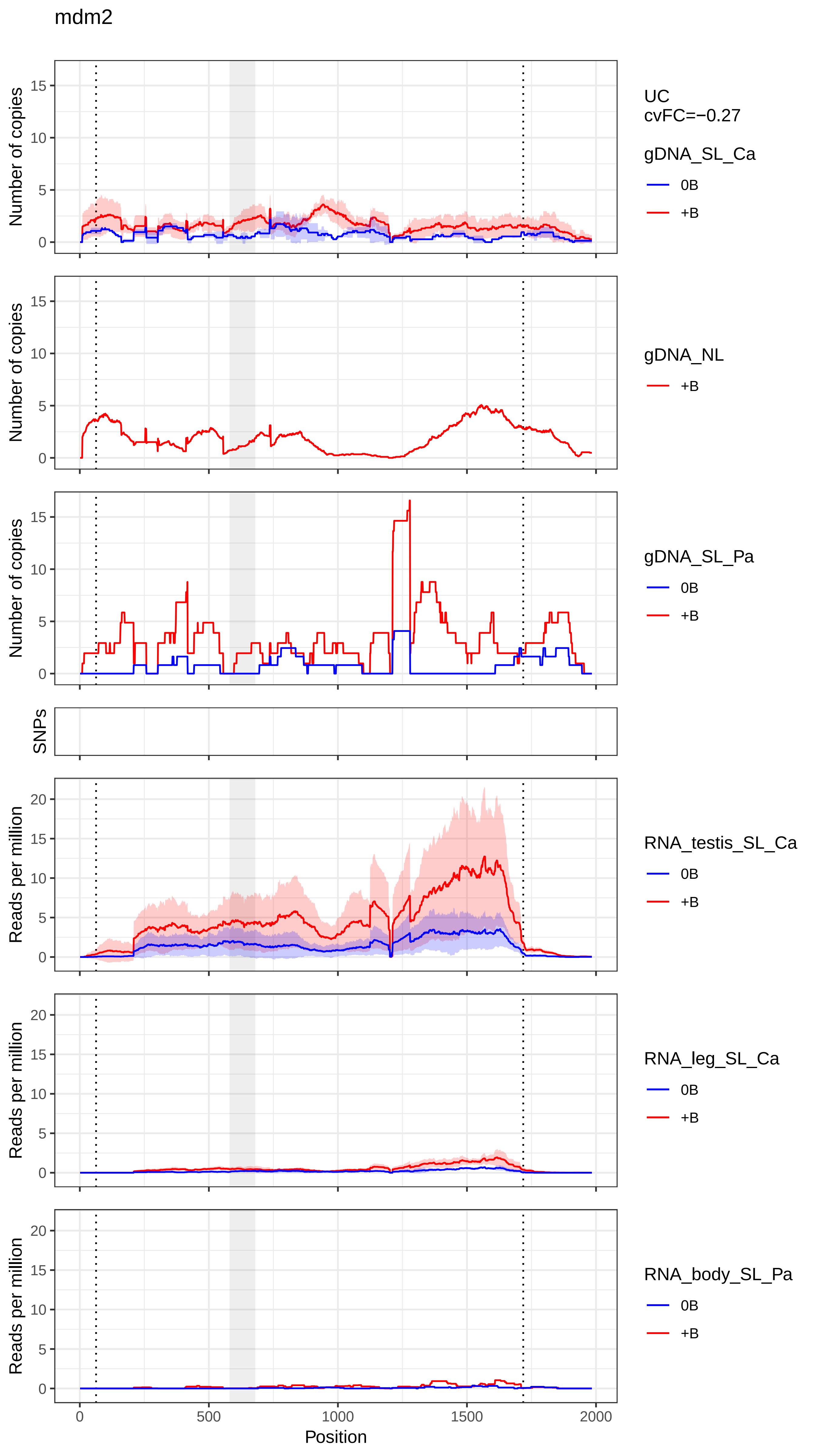

mdm2
Number of copies
UC
cvFC=−0.27
gDNA_SL_Ca
0B
+B
gDNA_NL
+B
gDNA_SL_Pa
0B
+B
SNPs
Reads per million
RNA_testis_SL_Ca
0B
+B
RNA_leg_SL_Ca
0B
+B
RNA_body_SL_Pa
0B
+B
0
500
1000
1500
2000
Position

nach

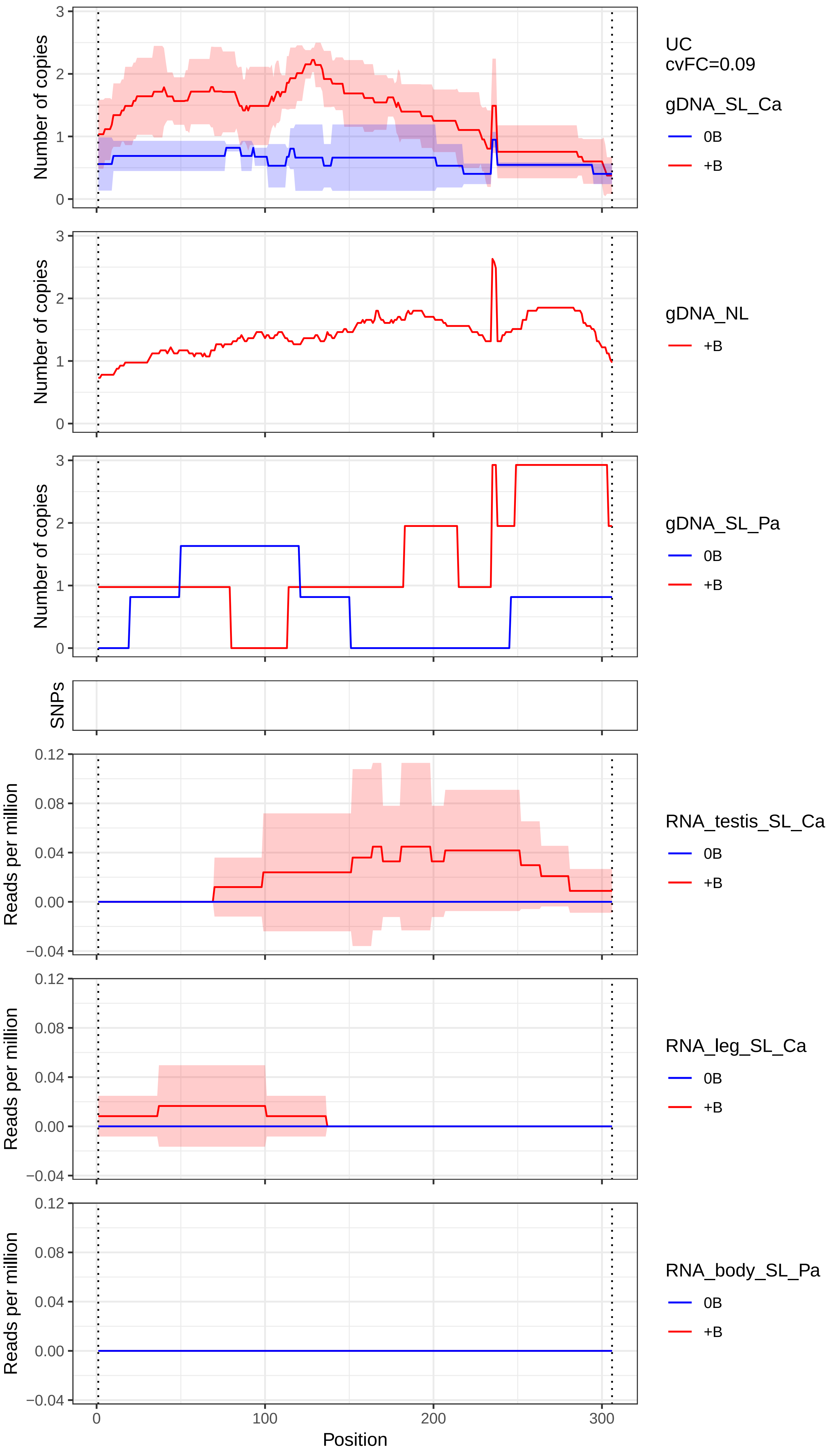

Number of copies
3
2
1
0
UC
cvFC=0.09
gDNA_SL_Ca
0B
+B
Number of copies
gDNA_NL
+B
Number of copies
gDNA_SL_Pa
0B
+B
SNPs
Reads per million
0.12
0.08
0.04
0.00
−0.04
RNA_testis_SL_Ca
0B
+B
Reads per million
RNA_leg_SL_Ca
0B
+B
Reads per million
RNA_body_SL_Pa
0B
+B
0
100
200
300
Position

nrc2a

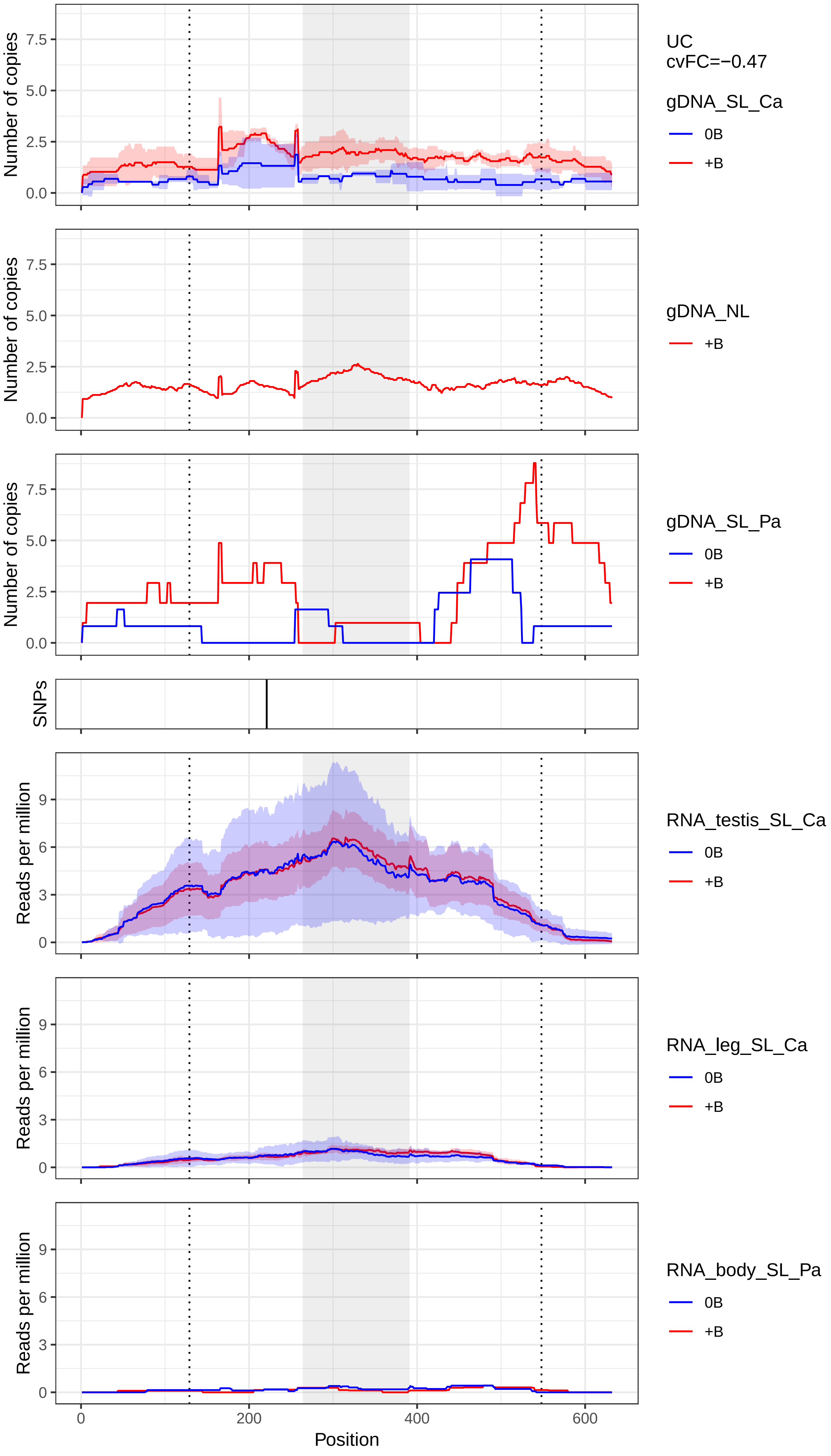

UC
cvFC=−0.47
gDNA_SL_Ca
0B
+B
gDNA_NL
+B
gDNA_SL_Pa
0B
+B
SNPs
RNA_testis_SL_Ca
0B
+B
RNA_leg_SL_Ca
0B
+B
RNA_body_SL_Pa
0B
+B
Number of copies
Reads per million
0.0
2.5
5.0
7.5
0
3
6
9
0
200
400
600
Position

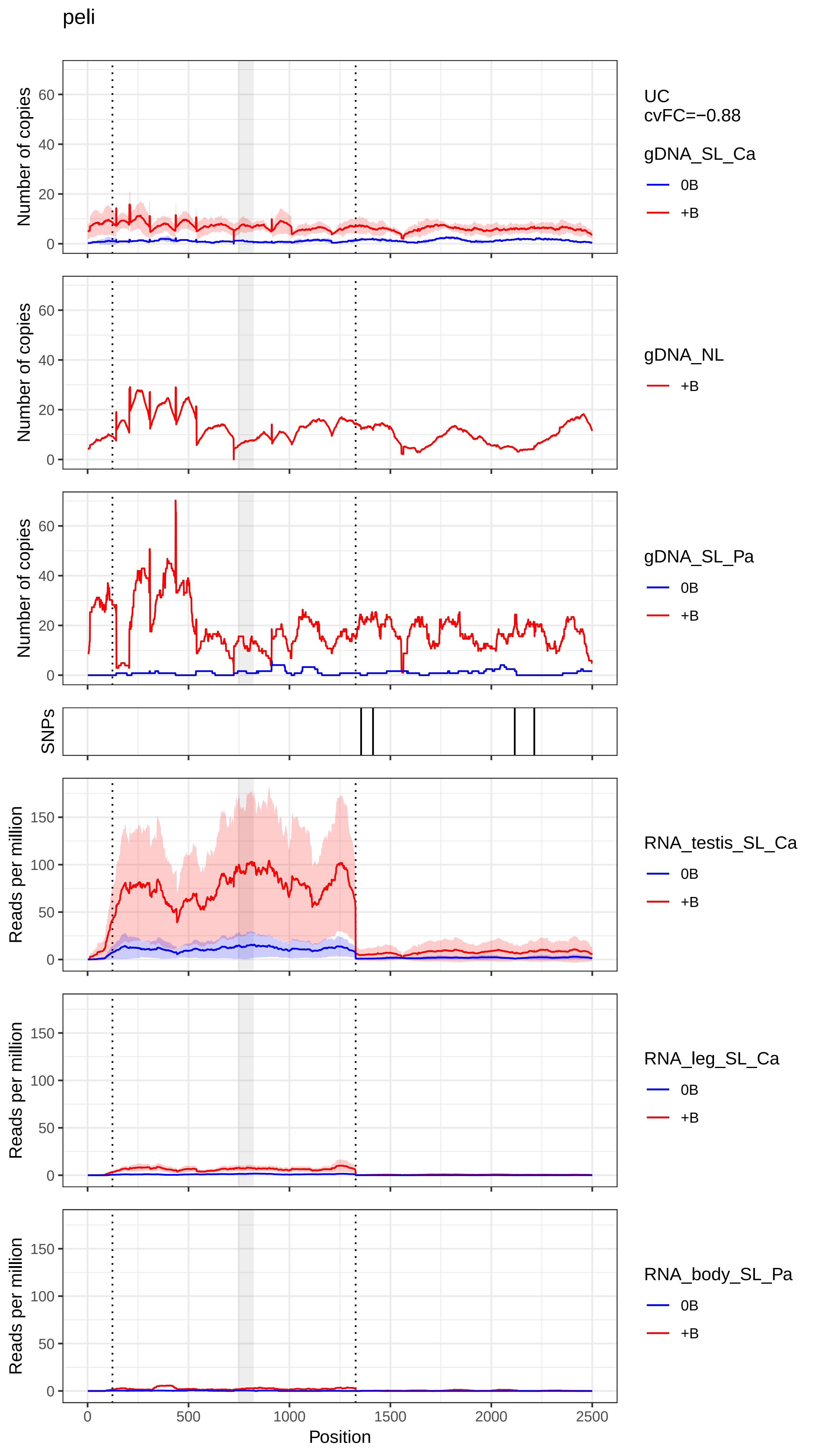
peli
UC
cvFC=−0.88
gDNA_SL_Ca
0B
+B
gDNA_NL
+B
gDNA_SL_Pa
0B
+B
SNPs
RNA_testis_SL_Ca
0B
+B
RNA_leg_SL_Ca
0B
+B
RNA_body_SL_Pa
0B
+B
Number of copies
Reads per million
Position
0
500
1000
1500
2000
2500
0
20
40
60
50
100
150

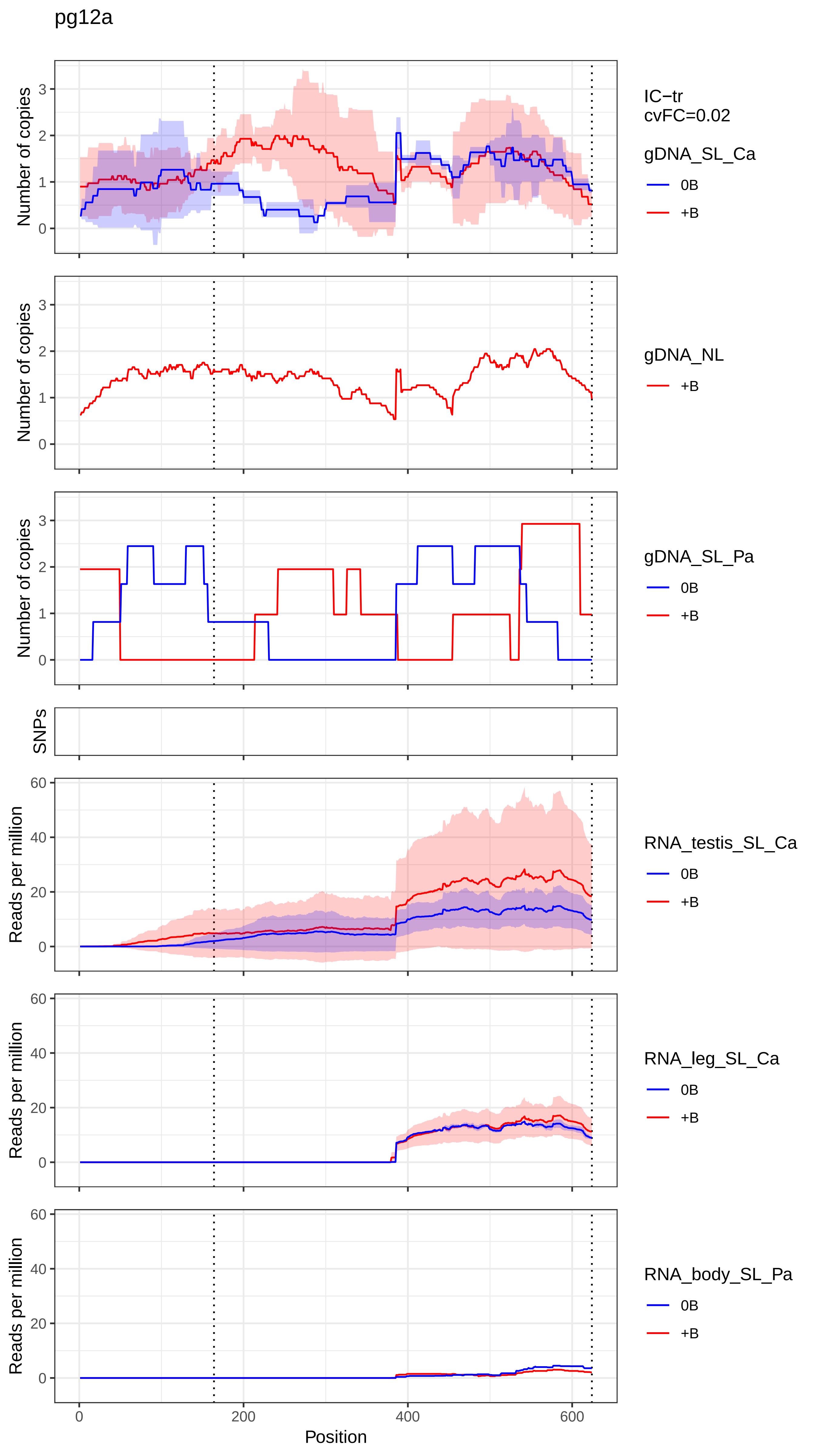

pg12a
IC−tr
cvFC=0.02
gDNA_SL_Ca
0B
+B
gDNA_NL
+B
gDNA_SL_Pa
0B
+B
SNPs
RNA_testis_SL_Ca
0B
+B
RNA_leg_SL_Ca
0B
+B
RNA_body_SL_Pa
0B
+B
Number of copies
Reads per million
Position
0
200
400
600

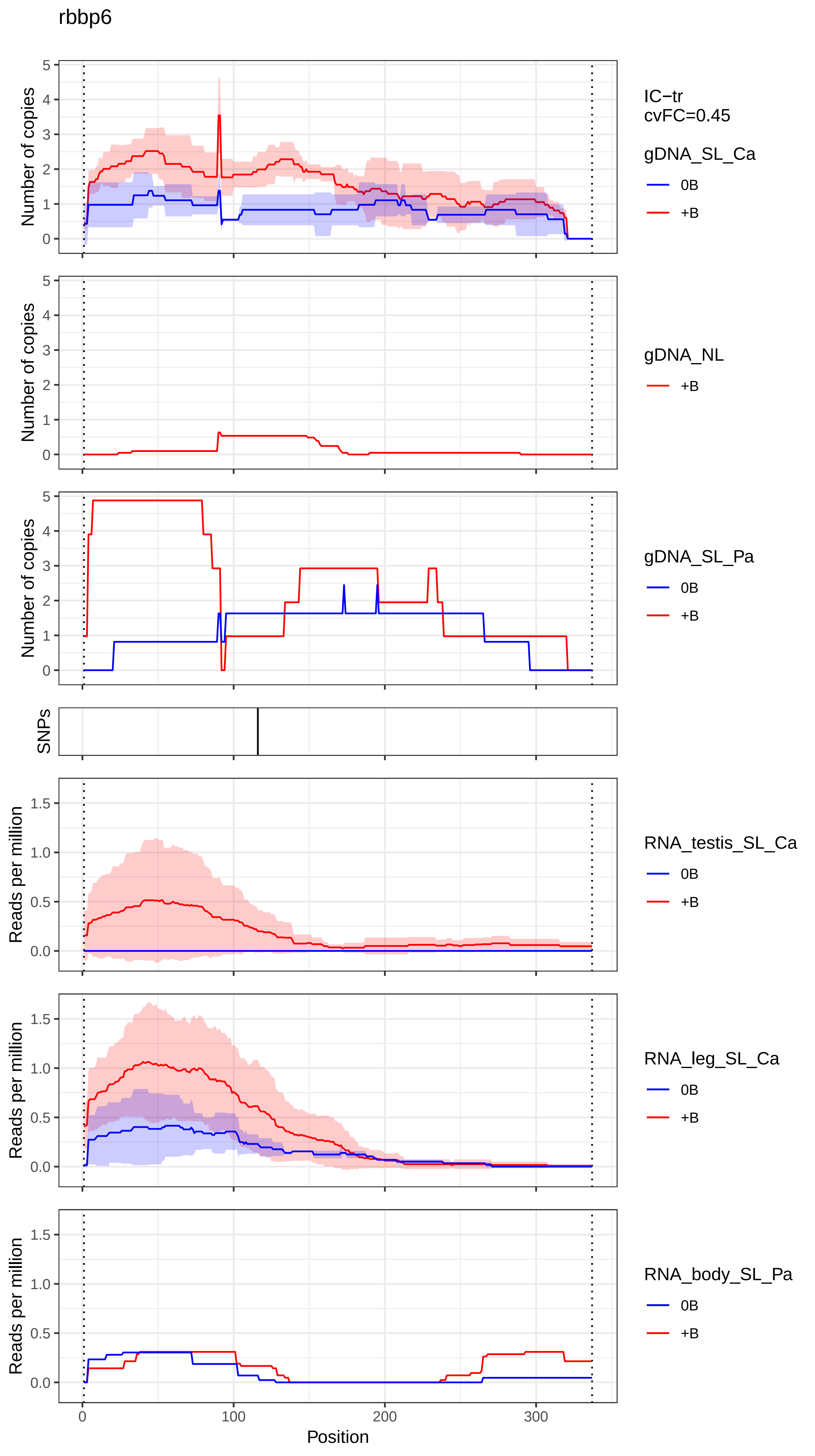
rbbp6
Number of copies
IC−tr
cvFC=0.45
gDNA_SL_Ca
0B
+B
gDNA_NL
+B
gDNA_SL_Pa
0B
+B
SNPs
Reads per million
RNA_testis_SL_Ca
0B
+B
RNA_leg_SL_Ca
0B
+B
RNA_body_SL_Pa
0B
+B
0
100
200
300
Position

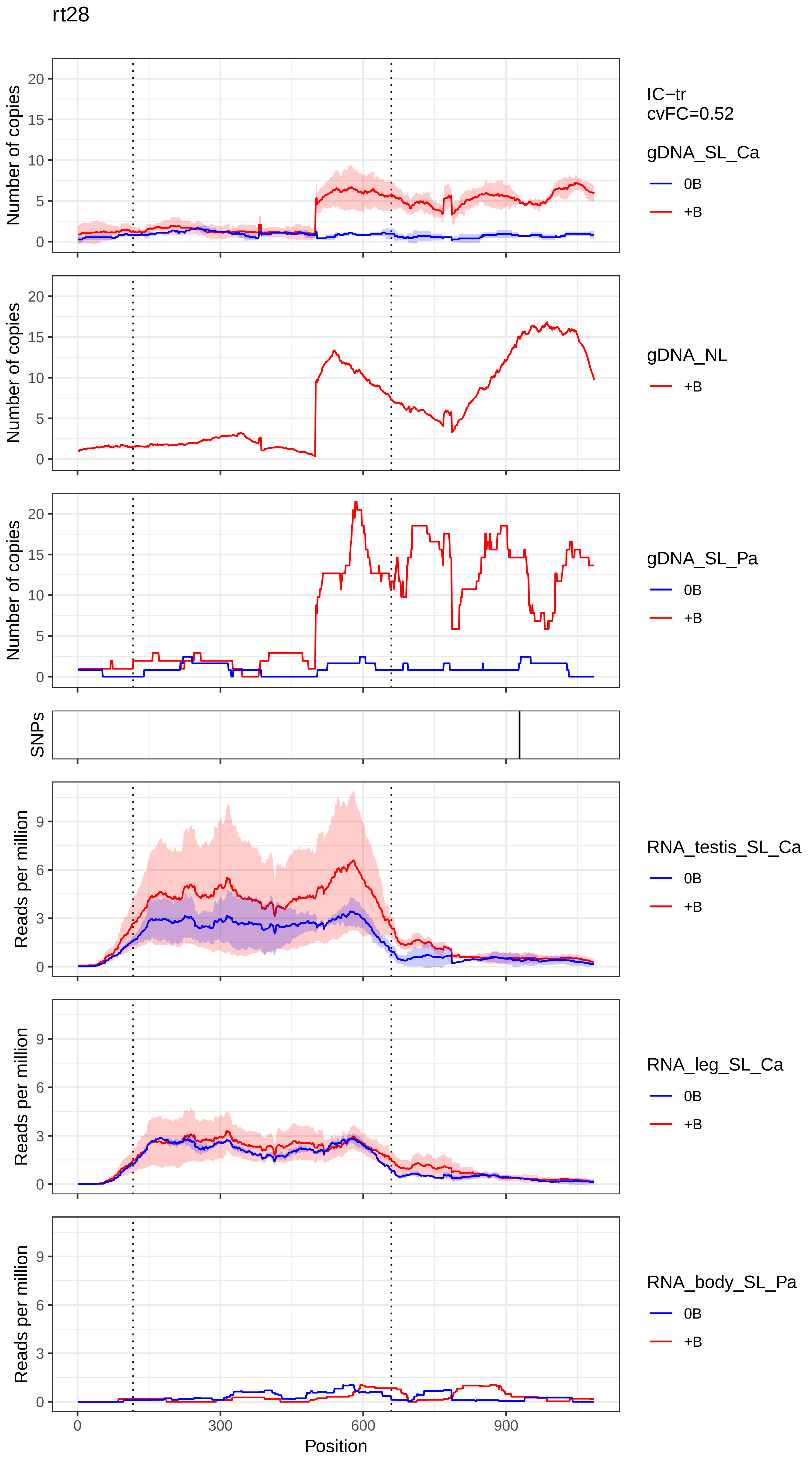
rt28
Number of copies
IC−tr
cvFC=0.52
gDNA_SL_Ca
0B
+B
gDNA_NL
+B
gDNA_SL_Pa
0B
+B
SNPs
Reads per million
RNA_testis_SL_Ca
0B
+B
RNA_leg_SL_Ca
0B
+B
RNA_body_SL_Pa
0B
+B
0
300
600
900
Position

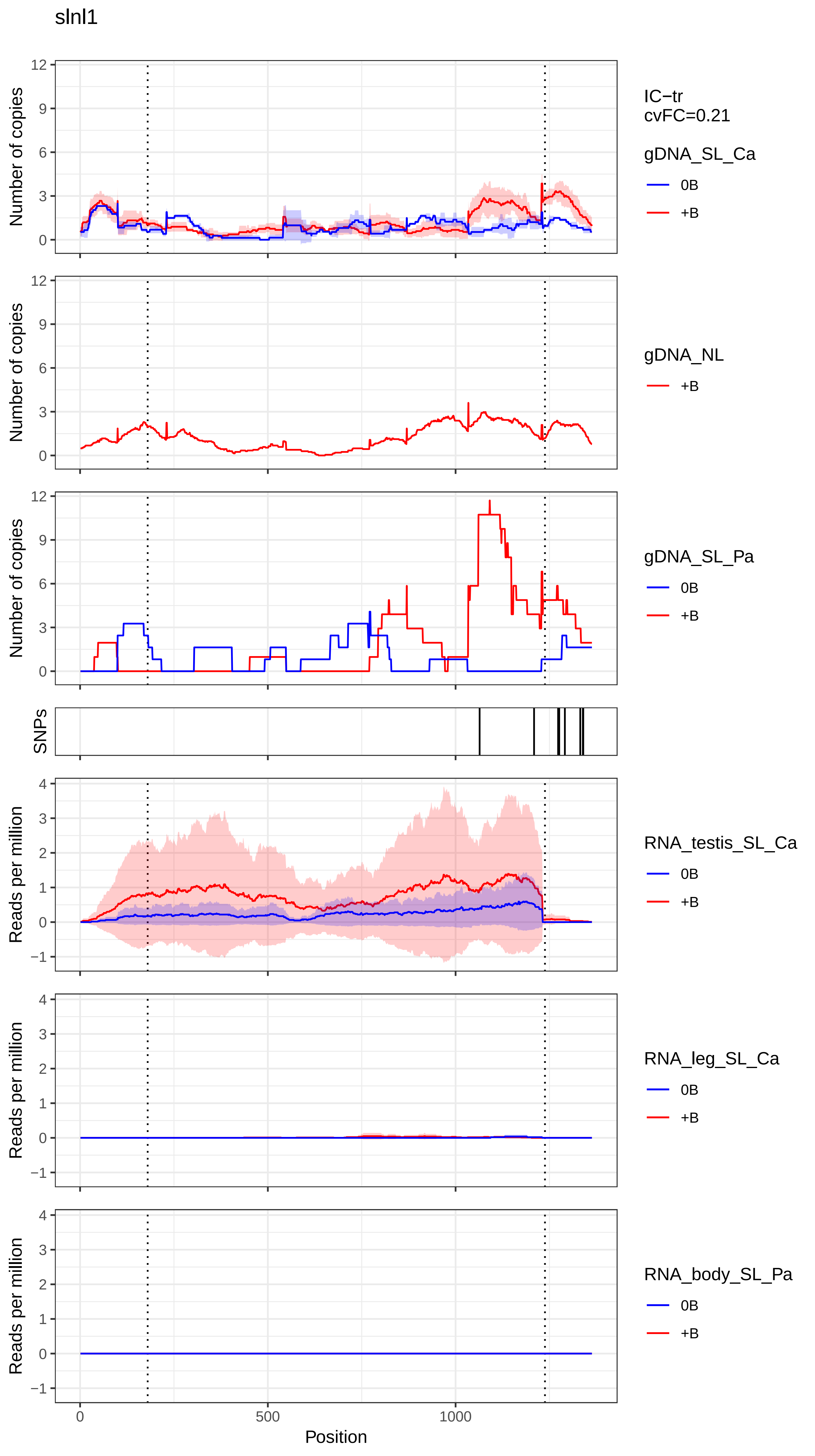
slnl1
Number of copies
IC−tr
cvFC=0.21
gDNA_SL_Ca
0B
+B
gDNA_NL
+B
gDNA_SL_Pa
0B
+B
SNPs
Reads per million
RNA_testis_SL_Ca
0B
+B
RNA_leg_SL_Ca
0B
+B
RNA_body_SL_Pa
0B
+B
0
500
1000
Position

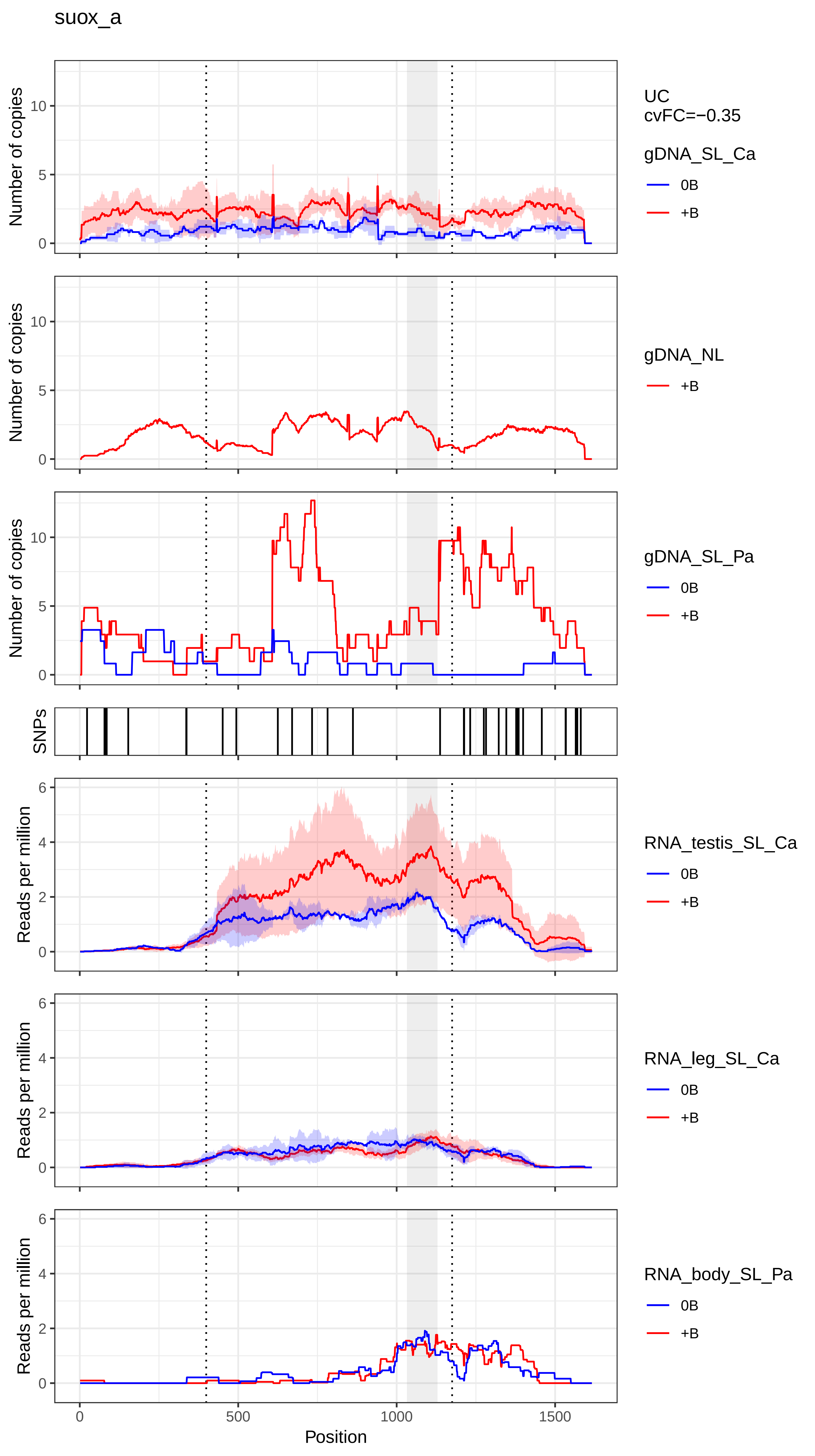
suox_a
Number of copies
UC
cvFC=−0.35
gDNA_SL_Ca
0B
+B
gDNA_NL
+B
gDNA_SL_Pa
0B
+B
SNPs
Reads per million
RNA_testis_SL_Ca
0B
+B
RNA_leg_SL_Ca
0B
+B
RNA_body_SL_Pa
0B
+B
0
500
1000
1500
Position

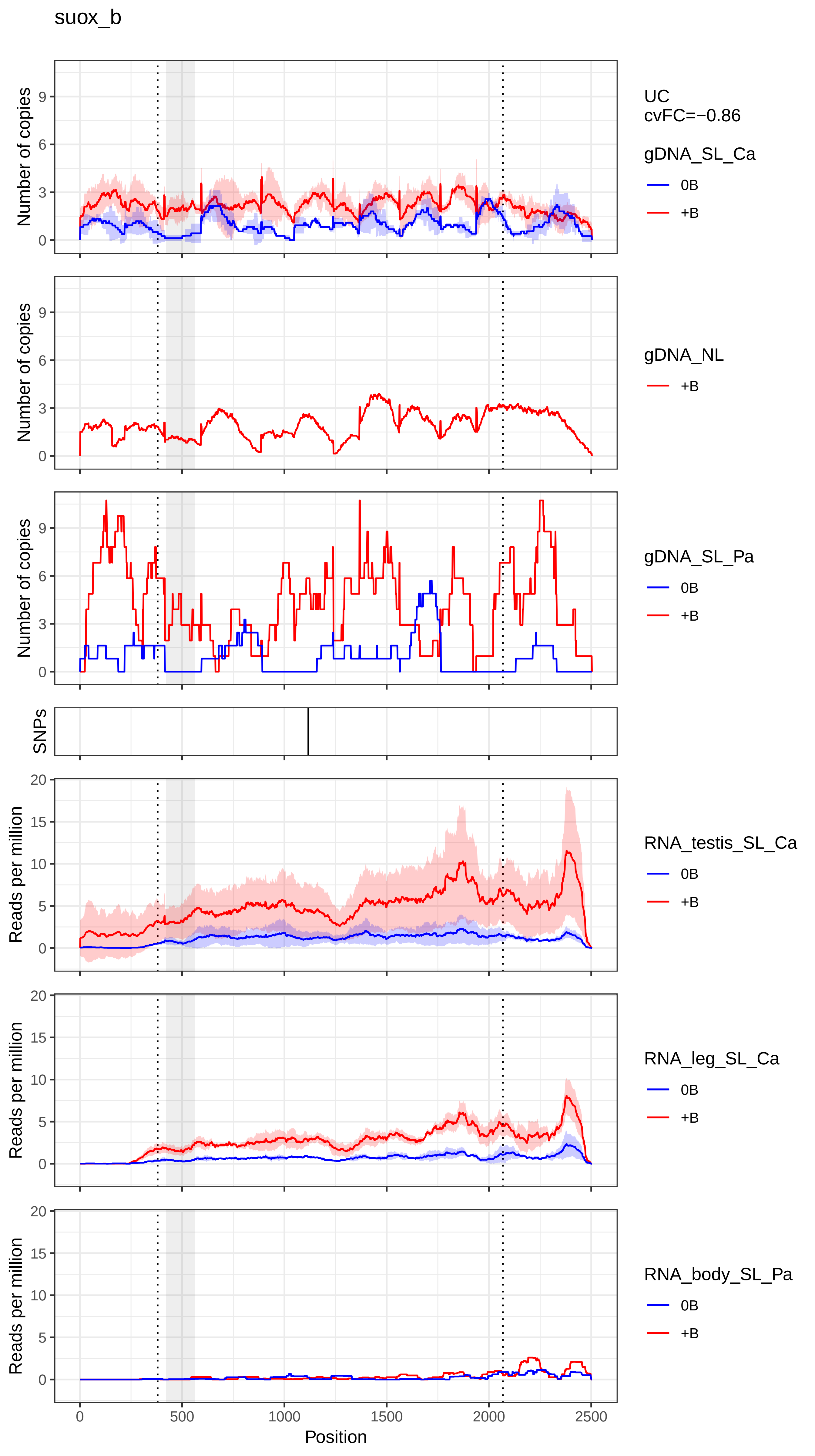

suox_b
UC
cvFC=−0.86
gDNA_SL_Ca
0B
+B
gDNA_NL
+B
gDNA_SL_Pa
0B
+B
SNPs
RNA_testis_SL_Ca
0B
+B
RNA_leg_SL_Ca
0B
+B
RNA_body_SL_Pa
0B
+B
Number of copies
Reads per million
Position
0
500
1000
1500
2000
2500

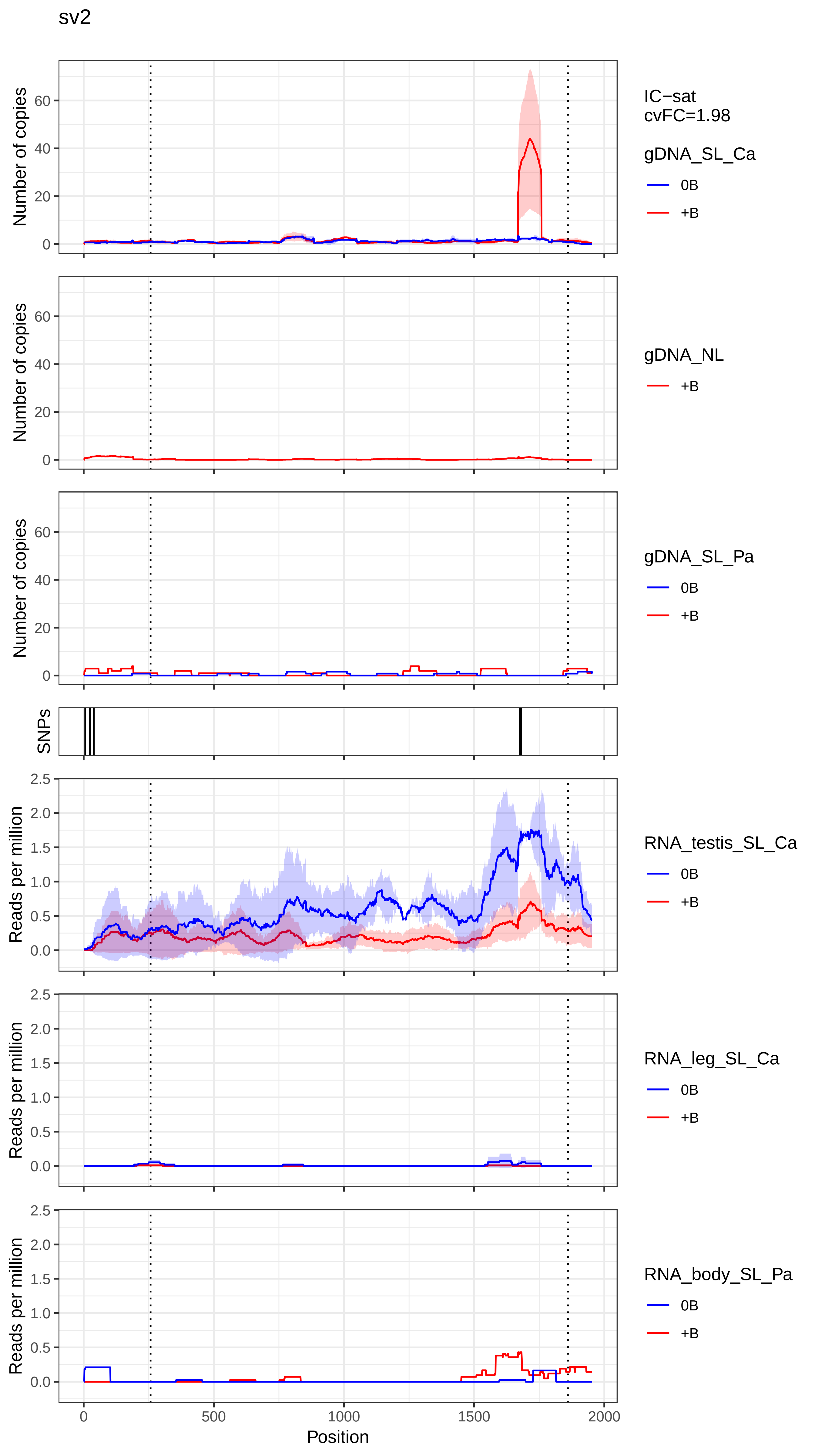

sv2
Number of copies
Number of copies
Number of copies
SNPs
Reads per million
Reads per million
Reads per million
Position
0
500
1000
1500
2000
IC−sat
cvFC=1.98
gDNA_SL_Ca
0B
+B
gDNA_NL
+B
gDNA_SL_Pa
0B
+B
RNA_testis_SL_Ca
0B
+B
RNA_leg_SL_Ca
0B
+B
RNA_body_SL_Pa
0B
+B

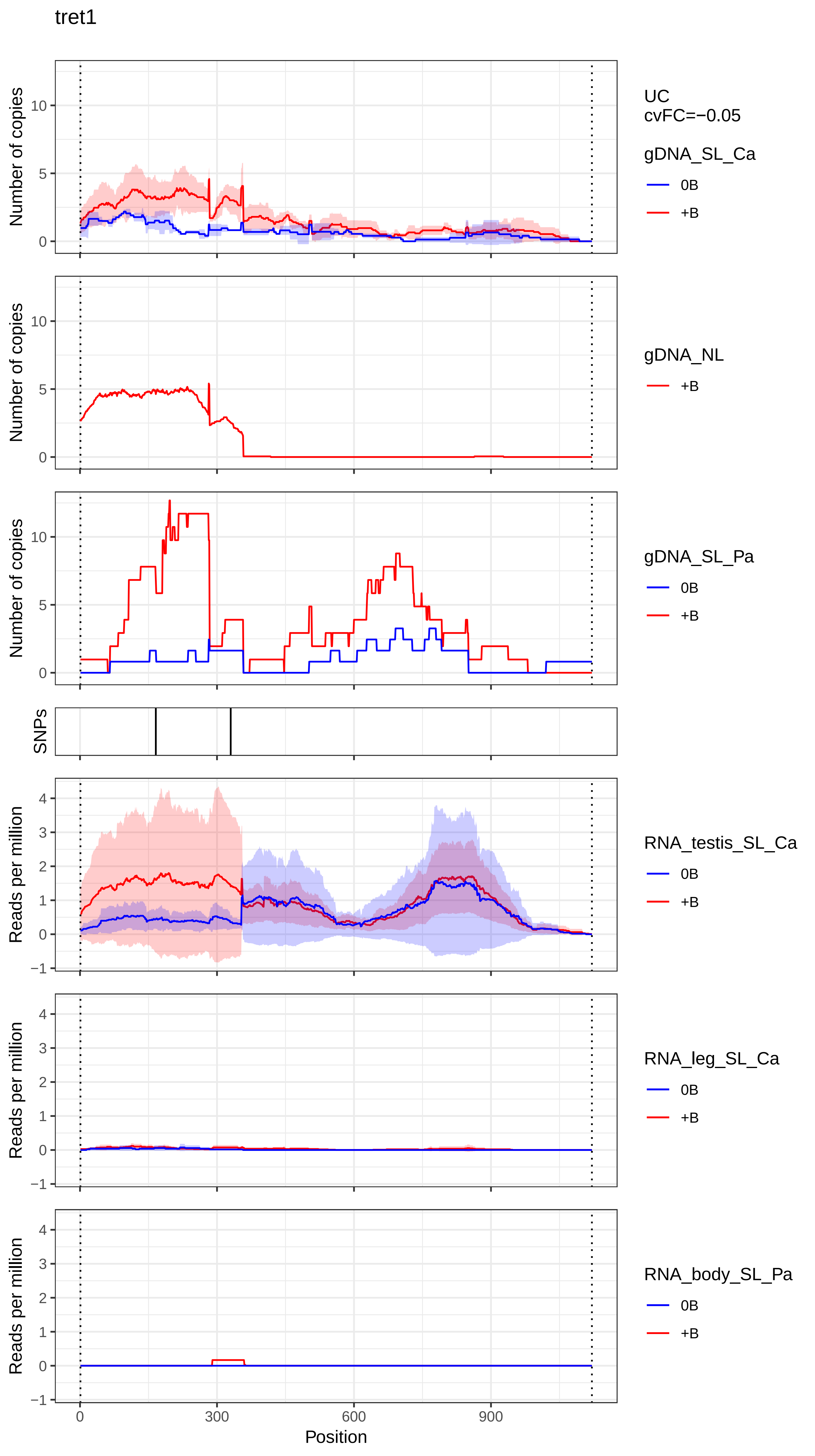

tret1
UC
cvFC=−0.05
gDNA_SL_Ca
0B
+B
gDNA_NL
+B
gDNA_SL_Pa
0B
+B
RNA_testis_SL_Ca
0B
+B
RNA_leg_SL_Ca
0B
+B
RNA_body_SL_Pa
0B
+B
Number of copies
SNPs
Reads per million
Position
0
300
600
900

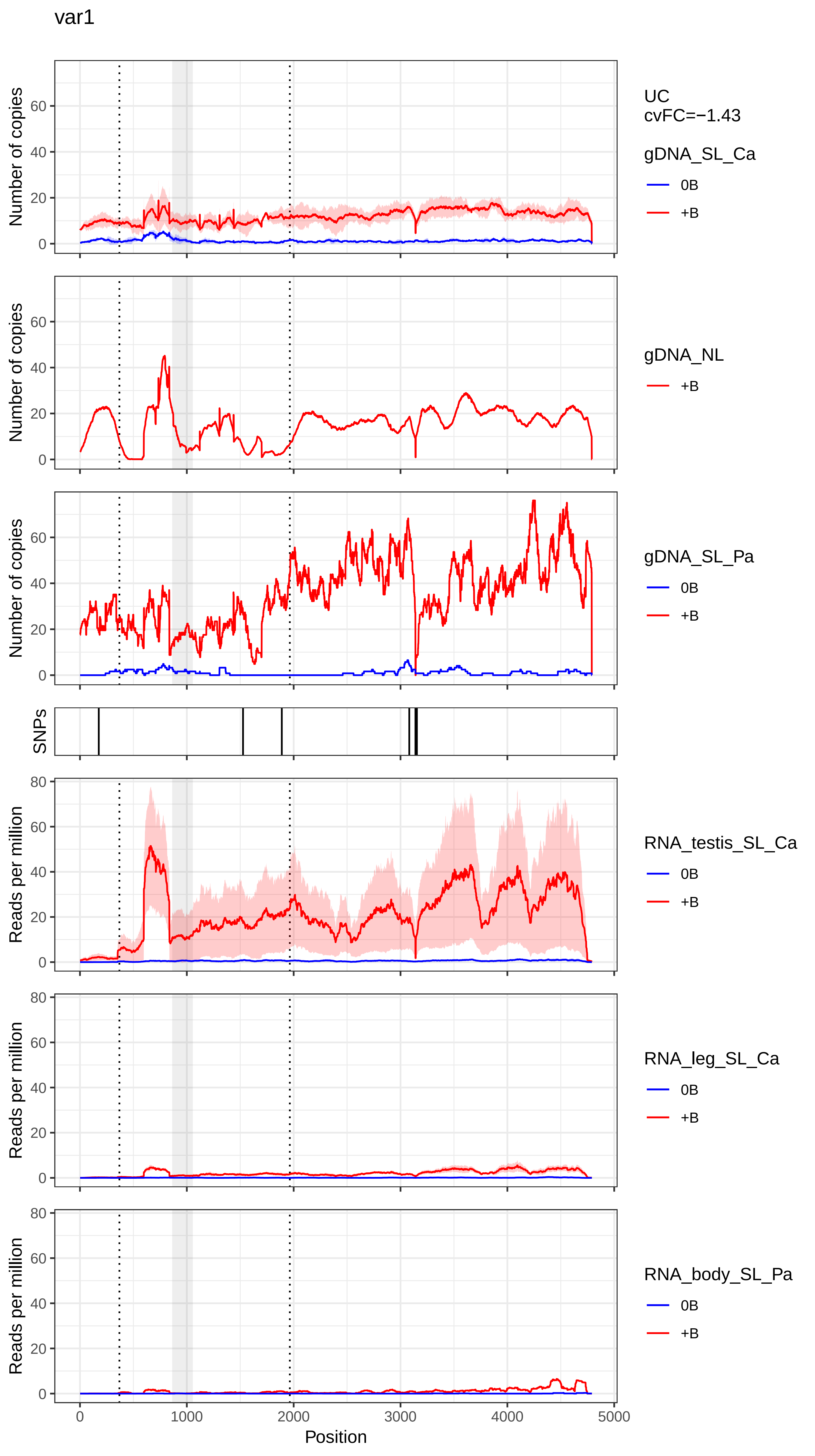
var1
UC
cvFC=−1.43
gDNA_SL_Ca
0B
+B
gDNA_NL
+B
gDNA_SL_Pa
0B
+B
SNPs
RNA_testis_SL_Ca
0B
+B
RNA_leg_SL_Ca
0B
+B
RNA_body_SL_Pa
0B
+B
Number of copies
Reads per million
Position
0
1000
2000
3000
4000
5000
0
20
40
60
80

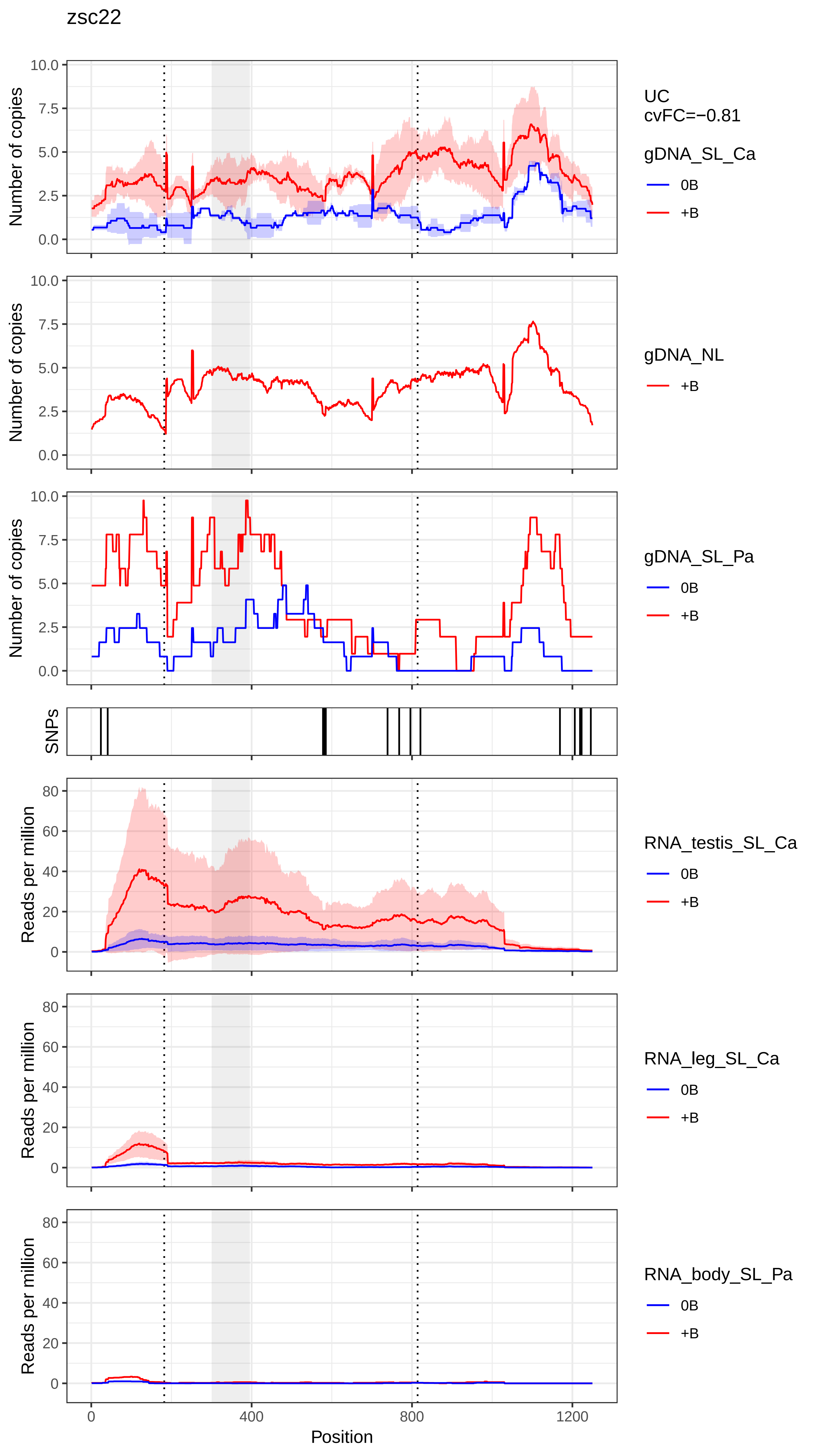
zsc22
UC
cvFC=−0.81
gDNA_SL_Ca
0B
+B
gDNA_NL
+B
gDNA_SL_Pa
0B
+B
SNPs
RNA_testis_SL_Ca
0B
+B
RNA_leg_SL_Ca
0B
+B
RNA_body_SL_Pa
0B
+B
Number of copies
Reads per million
Position
0
400
800
1200
